# When does gene flow facilitate evolutionary rescue?

**DOI:** 10.1101/622142

**Authors:** Matteo Tomasini, Stephan Peischl

## Abstract

Experimental and theoretical studies have highlighted the impact of gene flow on the probability of evolutionary rescue in structured habitats. Mathematical modelling and simulations of evolutionary rescue in spatially or otherwise structured populations showed that intermediate migration rates can often maximize the probability of rescue in gradually or abruptly deteriorating habitats. These theoretical results corroborate the positive effect of gene flow on evolutionary rescue that has been identified in experimental yeast populations. The observations that gene flow can facilitate adaptation are in seeming conflict with traditional population genetics results that show that gene flow usually hampers (local) adaptation. Identifying conditions for when gene flow facilitates survival chances of populations rather than reducing them remains a key unresolved theoretical question. We here present a simple analytically tractable model for evolutionary rescue in a two-deme model with gene flow. Our main result is a simple condition for when migration facilitates evolutionary rescue, as opposed as no migration. We further investigate the roles of asymmetries in gene flow and / or carrying capacities, and the effects of density regulation and local growth rates on evolutionary rescue.

## Introduction

Evolutionary rescue refers to the process of rapid adaptation to prevent extinction in the face of severe environmental change [Gomulkiewicz and Holt, 1995]. It is of particular interest in light of recent environmental and climatic change, with the potential to lead to new conservation strategies [Ashley et al., 2003]. Evolutionary rescue also plays a major role in other fields of public importance, such as the evolution of antibiotic or other treatment resistance (e.g. Normark and Normark [2002]), or resistance to pesticides (e.g. Chevillon et al. [1999]). Better understanding of evolutionary rescue is therefore critical in the context of global climatic change as well as in the field of evolutionary medicine. Experimental evolution studies of evolutionary rescue and antibiotic resistance are burgeoning (reviewed in Bell [2017]), empirical evidence for rescue under anthropogenic stress is now abundant [Hughes and Andersson, 2017, Bell, 2017], whereas evidence for rescue under natural conditions is difficult to obtain and more scarce (but see Vander Wal et al. [2013]).

The theoretical foundations for evolutionary rescue in single panmictic populations are laid out [Orr and Unckless, 2014] and several demographic genetic and extrinsic features that affect the chance for rescue have been identified (see table 1 in Carlson et al. [2014] for an overview), including the effects of recombination [Uecker and Hermisson, 2016], mating system [Uecker, 2017], intra-specific competition [Osmond and de Mazancourt, 2013, Bono et al., 2015], inter-specific competition [De Mazancourt et al., 2008], and phenotypic plasticity [Chevin et al., 2013, Carja and Plotkin, 2019]. A major goal of evolutionary rescue theory is to predict a population’s chance of survival in the face of severe stress. Key theoretical predictions of evolutionary rescue have been strikingly confirmed in laboratory conditions [Carlson et al., 2014], for instance using yeast populations exposed to high salt concentrations [Bell, 2013]. In particular, it was found that only sufficiently large populations could be expected to persist through adaptation [Lynch, 1993, Bell and Gonzalez, 2009, Samani and Bell, 2010, Bell and Gonzalez, 2011, Ramsayer et al., 2013, Bell, 2013]). A second feature that has been shown to facilitate the chance for evolutionary rescue theoretically as well as experimentally is standing genetic variation [Barrett and Schluter, 2008, Agashe et al., 2011, Lachapelle and Bell, 2012, Vander Wal et al., 2013, Ramsayer et al., 2013]. Despite these advances, however, predicting evolutionary outcomes outside of the lab remains extremely difficult [Gomulkiewicz and Shaw, 2013].

**Table 1:**
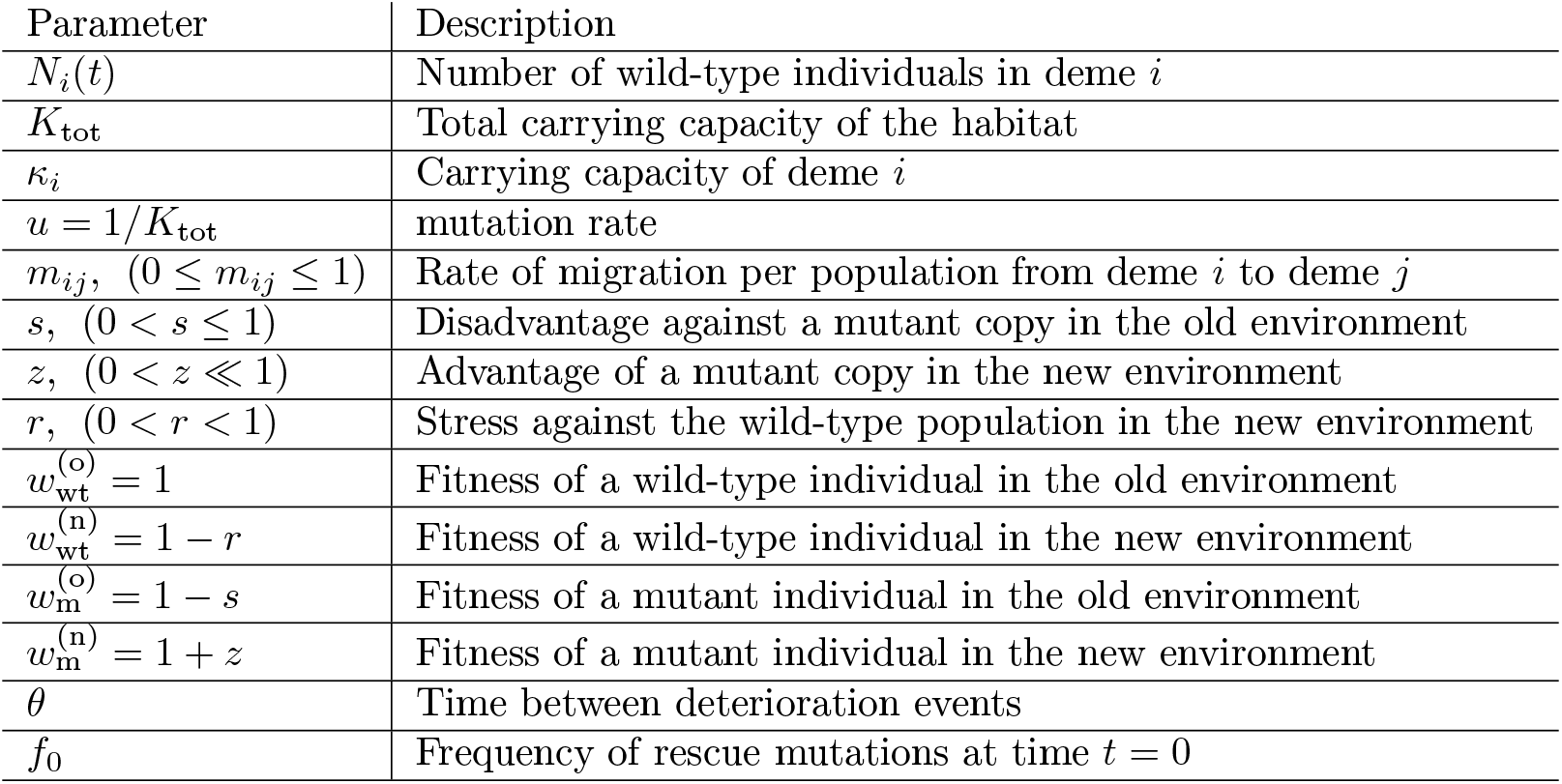
List and description of all parameters.

Evolutionary dynamics in spatially (or otherwise) structured populations can differ dramatically from those in well-mixed populations [Lion et al., 2011] and unexpected rescue mechanisms may arise in such settings [Peischl and Gilbert, 2018]. Empirical and experimental results have highlighted the importance of dispersal for evolutionary rescue in metapopulations subject to gradual environmental change. Using an experimental metapopulation of yeast exposed to gradually increasing environmental stress, Bell and Gonzalez [2011] showed that gene flow between different habitats can have positive effects on survival in changing environments, depending on dispersal distances and the speed of the environmental change. A detailed theoretical study of evolutionary rescue in structured populations using mathematical analysis and simulations confirmed that intermediate gene flow between populations can maximize the chance of rescue as compared to a population without gene flow [Uecker et al., 2014] in some cases. Uecker et al. [2014] identified two direct consequences of dispersal: (i) the unperturbed environment acts as a source for wild-type individuals that might mutate, thus increasing the chances of rescue, and (ii) dispersal moves mutant individuals to regions of the environment where the presence of the mutation is costly, leading to a net reduction of the mutant growth rate, and consequent lower rates of survival. The interplay between these two effects can often lead to situations in which the probability of rescue is maximized for an intermediate migration rate [Uecker et al., 2014]. In a continuous space model where the environment changes gradually across space and/or time, increased dispersal generally decreases the probability of establishment of rescue mutations, but it increases the effective population size of individuals that can contribute to evolutionary rescue [Kirkpatrick and Peischl, 2013]. Individual based simulations of gradually changing conditions and divergent selection between two habitats identified interactions of evolutionary rescue and local adaptation in a two-deme model [Bourne et al., 2014]. These results suggest that gene flow is beneficial for population survival only when divergent selection is relatively weak. These results were largely confirmed in a simulation study of a 2D metapopulation [Schiffers et al., 2013].

Although both theoretical and experimental studies have identified potentially positive effects of gene flow on survival in metapopulation models of evolutionary rescue, the exact conditions when gene flow is detrimental to survival and when not remain unclear. For instance, the observation that gene flow can facilitate rescue in a changing environment is in seeming conflict with more traditional results that show that dispersal does generally not have a positive effect on (local) adaptation [Bulmer, 1972, Holt and Gomulkiewicz, 1997, Lenormand, 2002]). High migration rates can lead to gene swamping in models with divergent selection pressures between different regions [Bulmer, 1972, Lenormand, 2002], thus reducing chances of survival during environmental change. Identifying conditions under which dispersal facilitates evolutionary rescue in spatially or otherwise structured populations remains a key unresolved question, both theoretically and empirically. In this article, we present an analytically tractable model with two demes that exchange migrants, and with temporal change in environmental conditions. We focus on the case where the two demes deteriorate at different points in time, such that gene flow between the populations influences both the demographic as well as the evolutionary dynamics of evolutionary rescue. In the new environmental conditions, growth rates are negative and the population faces eventual extinction. We consider rescue mutations at a single locus and assume that they are counter-selected in the original environmental conditions. We derive conditions for when gene flow facilitates evolutionary rescue as compared to two populations without gene flow. We study the role of asymmetric migration rates or asymmetric carrying capacities (both cases can lead to source-sink dynamics, see Holt [1985], Pulliam [1988]), study the contributions of de novo mutations vs. standing genetic variation, and investigate the role of local growth rates and density regulation within demes. Our aim is to understand when gene flow facilitates evolutionary rescue, and to disentangle the interactions between the strength of selection for rescue mutations, the speed and severity of environmental change, and the amount and mode of dispersal.

## Model

We consider a haploid population with discrete non-overlapping generations, subdivided into two demes, labeled 1 and 2, with gene flow between them. Individuals migrate from deme *i* to deme *j* with probability *m_ij_* (*i, j* ∈ {1, 2}). Fitness is determined by a single locus with two alleles: a wild-type allele and a mutant allele. We distinguish two possible environmental states. At the beginning both demes are in what we call the non-deteriorated state (or “old” state) and are at demographic equilibrium, filled with *κ_i_* individuals. The total population size is therefore *K*_tot_ = *κ*_1_+*κ*_2_. At time *t* = 0 deme 1 deteriorates (that is, it is now in the “new” state). In the deteriorated environment, wild-type individuals have absolute fitness 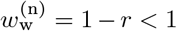, such that the population size in deme 1 declines at rate *r*. After *θ* generations, deme 2 deteriorates too and local population size starts to decline at the same rate as in deme 1. In the absence of adaptation to the novel environmental conditions both demes will eventually go extinct. We assume that rescue mutations that restore positive growth rates in the new environment occur at rate *u* per individual and generation, and we ignore back mutations. The absolute fitness of a mutant individual is 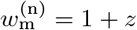 in the new habitat (*z* > 0). We assume that the mutation is detrimental in the old environment and denote its carriers fitness by 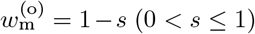. We call *r* the environmental stress due to deterioration, and *s* and *z* are the selection coefficients of the mutant allele in the old and new state, respectively. We will call “phase 1” the phase in which the two demes have different environments (0 *< t < θ*) and “phase 2” the phase in which both demes are deteriorated. See table 1 for a description of all the parameters of the model.

### Probability of rescue

Let *P*_rescue_ denote the probability that a rescue mutation occurs and escapes genetic drift, such that it will increase in frequency and eventually restore a positive growth rate and rescue the population from extinction. To calculate the probability of rescue, one needs to take into account two ingredients: (i) the number of mutations entering the population in each generation and (ii) the probability of establishment of each single mutant copy in the population. In a single population, one can write the probability of rescue as

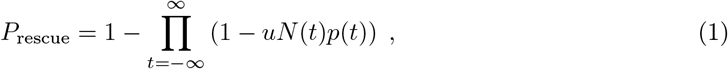

where *uN* (*t*) is the expected number of mutations entering the population in each generation, and *p*(*t*) is the probability that the mutation establishes and rescues the population [e.g., Gomulkiewicz and Holt, 1995]. We consider times from −∞ to +∞ here for mathematical convenience. Rescue mutations have a negligible probability of permanent establishment if they occur too early (at negative times *t* ≪ 0), as they are deleterious everywhere before phase 1. Similarly, for large times (*t* ≫ 0), the population will be extinct if no rescue mutation was successful before that.

Evolutionary rescue can stem from standing genetic variation, with probability *P*_sgv_, or from *de novo* mutations, with probability *P*_dn_. We define *de novo* mutations as mutations that arose after the first deterioration event occurred (that is, after time *t* = 0). We can thus write:

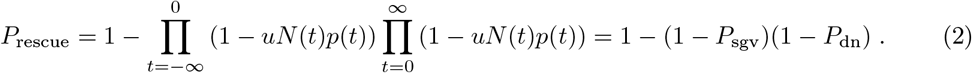

Mutations that occur before phase 2 (that is, that occur before all demes are deteriorated) have different probabilities of establishment *p*^(1)^(*t*) and *p*^(2)^(*t*) depending on the deme in which they occur and the time at which they occur. However, currently no analytic solution is known for the establishment probabilities in this case. To proceed further we ignore the temporal heterogeneity in fitness values and use the current environmental conditions to calculate establishment probabilities using the results from Tomasini and Peischl [2018] for a time-homogeneous two-deme model (assuming a large population size and small selection coefficient, i.e., 1*/N < z* ≪ 1). This should be a good approximation if *θ* ≫ 0, since the fate of mutations in temporally changing environments is determined in the first few generations after they occur [Peischl and Kirkpatrick, 2012] and the contribution of mutations occurring just before environments change will be negligible. In contrast, if *θ* ≈ 0, the change in environmental conditions is almost instantaneous across all demes, such that population structure and migration would have virtually no effect on evolutionary rescue [Uecker et al., 2014]. During phase 2, when the two demes are in the same environmental state, the probability of establishment is simply 2*z* [Haldane, 1927]. Tomasini and Peischl [2018] use branching processes to obtain the probability of establishment of mutations under divergent selection, as is the case during phase 1. The expression is shown here for a case with symmetric migration (*m*_12_ = *m*_21_ = *m/*2) [Tomasini and Peischl, 2018]. In the symmetric case, we define the rate of migration from one deme to the other as *m/*2 for consistency with the island model with *D* demes [Uecker et al., 2014], where *m_ij_* = *m/D*, for *i, j* ∈ {1, … *D*}. The probabilities of establishment for the two-deme model with symmetric migration are:

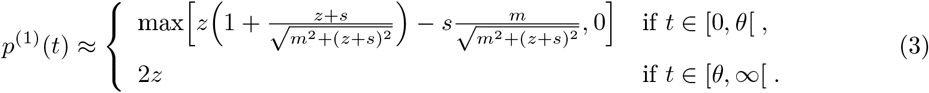

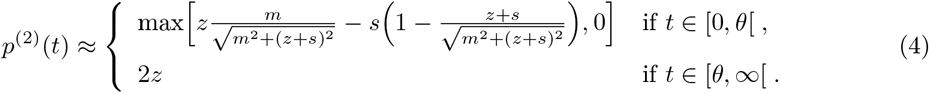

Because mutations have a negligible probability to establish at *t* ≪ 0 (see discussion before equation (2)), the probability of rescue due to standing genetic variation, *P*_sgv_, can be calculated as the probability of establishment of the mutations present in the population at time *t* = 0 due to mutation-selection balance. We can then write

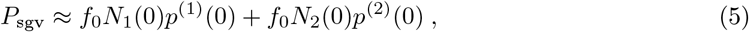

where *f*_0_ is the frequency of rescue mutations in each of the demes at time *t* = 0. Similarly, the total probability due to *de novo* mutations is given by

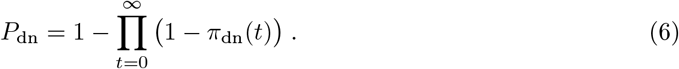

where we approximate the joint probability that a copy of the rescue mutation will occur in generation *t* and then establish permanently by

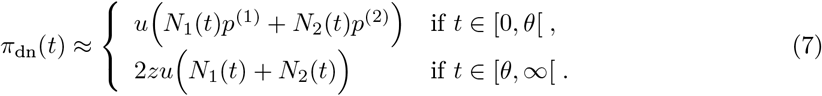

To simplify calculations, we use that 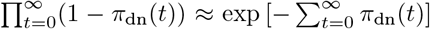 if *π*_dn_ is small, and for further simplicity, we do the calculation in continuous time, so that we can switch the sum for an integral. The probability of rescue from *de novo* mutations is then

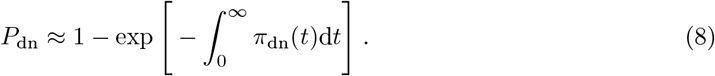

### Population dynamics

In order to calculate (6) and (7), we need to explicitly calculate the wild-type population sizes *N*_1_(*t*) and *N*_2_(*t*) for *t* ≥ 0. We assume that mutants are rare and hence we do not explicitly model their influence on demography. The only case where the number of mutants is large enough to effectively play a role is when a mutation is already on its way to establishment. We model the population dynamics as continuous in time, as we did in (8), and further assume that the mutation rate is low and neglect the number of wild-type individuals lost due to mutation. We assume that population growth and density regulation keep population density in deme 2 at carrying capacity, that is *N*_2_(*t*) = *κ*_2_, during phase 1. Population size in deme 1 then follows the differential equation

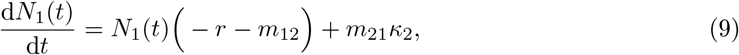

with initial condition *N*_1_(0) = *κ*_1_. During phase 2 (*t* ≥ *θ*), when both demes are deteriorated, *N*_1_(*t*) and *N*_2_(*t*) follow

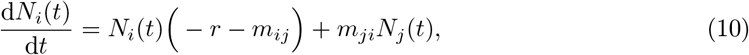

where *i, j* ∈ {1, 2} and *i* ≠ *j*. Solutions can be obtained straightforwardly – more details are given in the supplemental material (Appendix A, *e.g.* equation (S4) shows the solution for *i* = 1). Figure 1 shows the typical population dynamic trajectories during an evolutionary rescue event. In the absence of evolutionary rescue, population density would continue decaying until it reaches *N* = 0.

**Figure 1:**
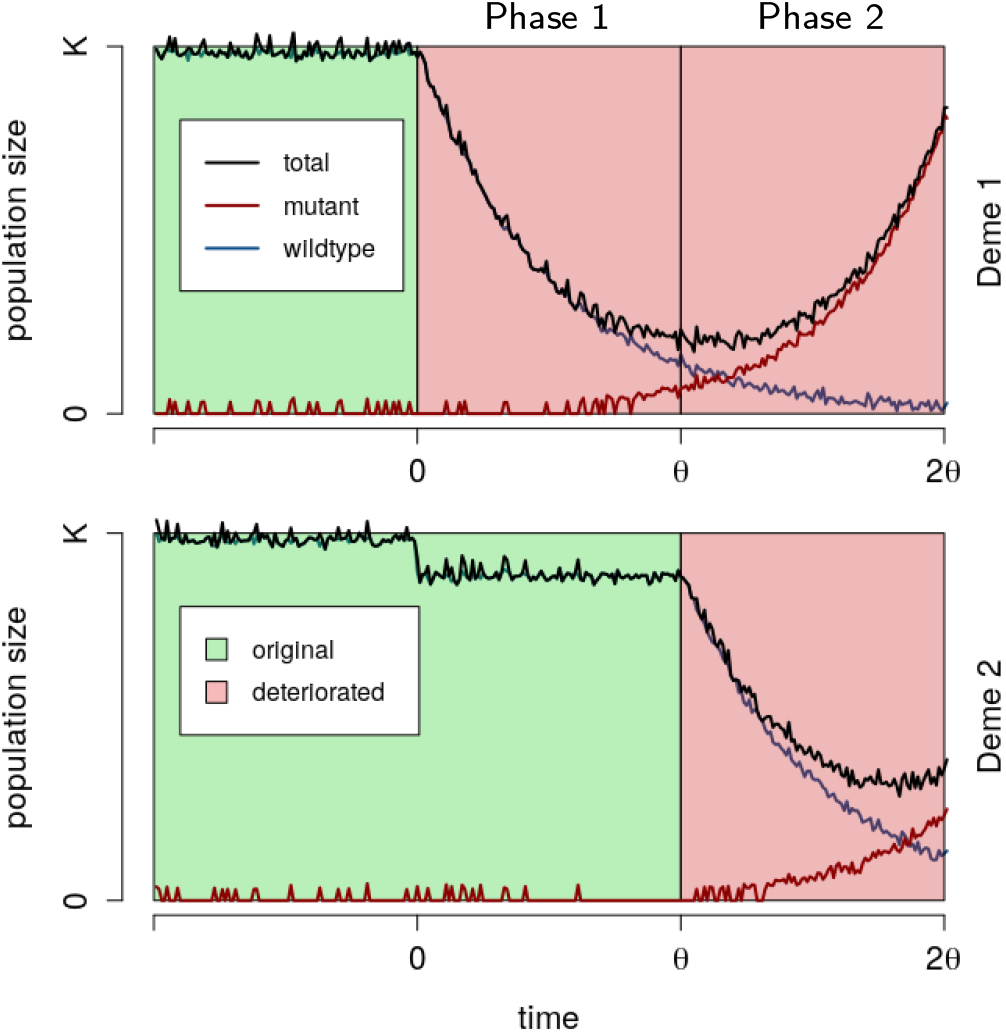
Schematic representation of evolutionary rescue in our model. On the upper panel, we show the population density in deme 1, in the lower panel the population density in deme 2. Deme 1 deteriorates at time *t* = 0, and deme 2 deteriorates at *t* = *θ*. The total count of individuals in deme 1 exhibits the typical “U-shape” associated with evolutionary rescue [Gomulkiewicz and Holt, 1995] (the same would be true in deme 2 if we extended the *x*-axis). In deme 2, in phase 1 we depict the number of individuals present just before density regulation. The drop in population observed during this phase depends on the migration rate.

### Simulation model

We performed stochastic simulations replicating biological processes to validate and extend our analytical findings. We filled a habitat with 20,000 individuals divided into two demes, labelled *i* = 1, 2, with carrying capacities *κ_i_*. We fixed the mutation rate at *u* = 1*/K*_tot_ = 5 × 10^*−*5^, so that in a non-deteriorated habitat at carrying capacity on average one new mutant enters the population per generation. Increasing (decreasing) *K*_tot_*u* will mainly lead to an increase (decrease) of the total rescue probability, and we hence keep *K*_tot_*u* fixed throughout the paper. The initial mutant frequency *f*_0_ was assumed at mutation-selection equilibrium, *f*_0_ = *u/s* [Gillespie, 2004]. At *t* = 0, deme 1 deteriorated, and at *t* = *θ* deme 2 deteriorated. Individuals in each deme reproduced, mutated and migrated, followed by density regulation. Generations are discrete and non-overlapping such that every generation the parental generation is replaced by its offspring. Each individual had Poisson distributed number of offspring with its mean proportional to the individuals fitness *w* (see table 1 for the definitions of fitnesses *w*). Every generation new mutants entered the population via binomial sampling from the wild-type population with probability *u*. Migration was also modeled as a binomial sampling from the local populations, where migrants from each deme *i* are sampled with probability *m_ij_* (*i, j* ∈ {1, 2}, *i* ≠ *j*). Density regulation was applied only to deme 2 when *t < θ* (non-deteriorated deme), and consisted in bringing the deme back to carrying capacity at the end of the generation. The genetic composition of the regulated deme was composed by binomial sampling, thus maintaining wild types and mutants in the non-perturbed deme at the same frequency that they reached after reproduction, mutation and migration. We run the simulation for two epochs of *θ* generations and add a burn-off period of 500 generations. Rescue was attained if at any moment during the simulation the number of mutants reaches *K*_tot_*/*2. We performed 2000 replicates for each parameter combination, and the probability of rescue is calculated as the proportion of replicates in which rescue occurred.

### Data availability

The source code for our simulations is available at the GitHub repository https://github.com/mtomasini/EvolutionaryRescue.

## Results

### Probability of rescue if mutations are lethal in the old environment

We start by evaluating (2) for the symmetric case where *κ*_1_ = *κ*_2_ = *κ* and *m*_12_ = *m*_21_ = *m/*2. Furthermore, we assume that the mutation is lethal in the old environment (*s* = 1), hence each rescue event will result from a *de novo* mutation. This allows us to outline our main results in a simple model and to provide some intuition about the involved mechanisms at play. We relax these assumptions later. Figure 2A shows the total probability of rescue (equation (2)) as a function of the migration rate, as well as the decomposition into mutations occurring during and after the deterioration of the environment. We observe that the probability of rescue with respect to migration is maximized for an intermediate migration rate for the parameter values used in Figure 1. This is consistent with previous results [Uecker et al., 2014]. The existence of an optimal intermediate migration rate reflects two effects that are at play here. On one hand the non-deteriorated deme acts as a source of wild-type individuals, preventing extinction in deme 1, thus increasing the chance for rescue to occur. On the other hand, too much migration between demes prevents rescue mutations from establishing despite being positively selected in one of the two demes, a process called gene swamping [Bulmer, 1972, Lenormand, 2002, Tomasini and Peischl, 2018] (Fig. 2, also see the discussion in the last section of Appendix A in the supplemental material). The limit beyond which gene flow causes swamping is *m > zs/*(*s* − *z*) (see red line in Fig. 2A) [Bulmer, 1972, Lenormand, 2002, Tomasini and Peischl, 2018]. Hence, for large migration rates, rescue can only occur during phase 2. In addition to these two processes, increasing the migration rate should also lead to an increased flux of individuals moving from deme 2 to deme 1, which would increase the total wild-type population size at the beginning of phase 2 (see supplemental material, Appendix A). Thus, we expect a mild positive effect on evolutionary rescue during phase 2 when increasing *m* (Fig. 2, also supplementary material, fig. S1). The mild positive effect of large migration during phase 2 stems from the fact that at time *t* = *θ* the number of individuals in deme 1, maintained exclusively by the influx of individuals from deme 2, increases with increasing migration rate (see supplemental material, Appendix A), the two demes behaving like one population. Because a larger population size increases the chance for rescue, our model predicts a slight increase of rescue for very large migration rates. This can be seen directly from equation (7).

**Figure 2:**
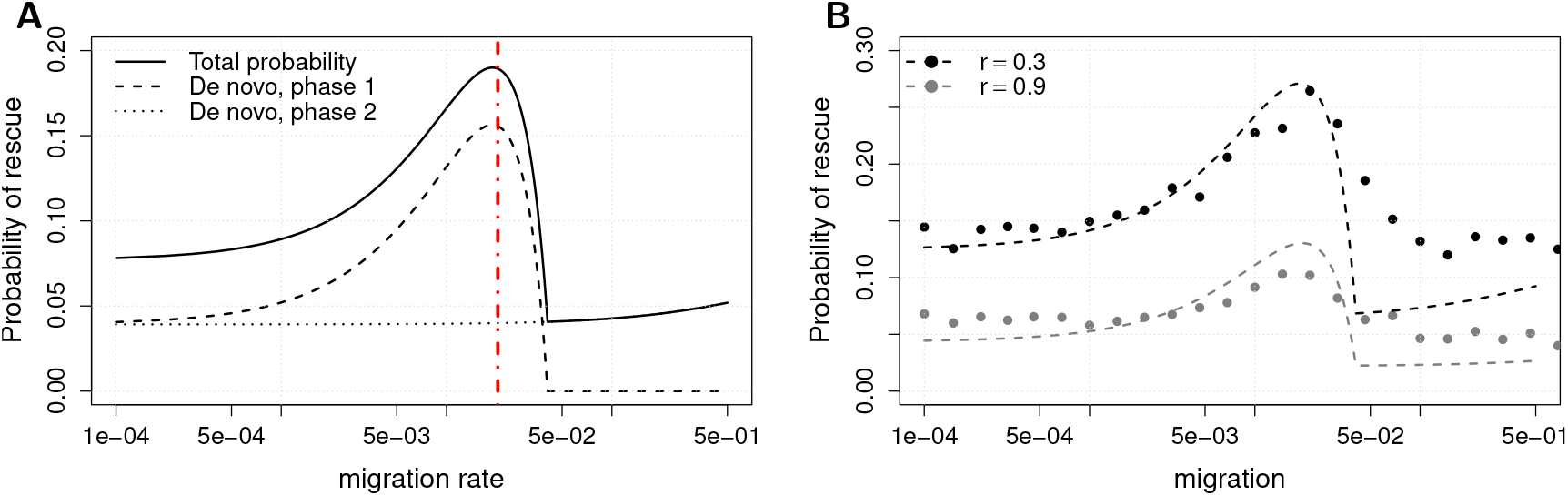
(A) The total probability of rescue and its decomposition in terms of *de novo* mutations during phases 1 and 2. The red vertical line represents the theoretical limit beyond which gene swamp disrupts rescue in phase 1. Parameters are *z* = 0.02, *s* = 1.0, *r* = 0.5 and *θ* = 500. (B) Comparison between simulations and prediction (equation 2), parameters are *z* = 0.02, *s* = 1.0 and *θ* = 500, in black *r* = 0.3 and in gray *r* = 0.9.

Figure 2B shows comparison with simulations and reveals a very good fit of our analytical approximation for low to intermediate migration rates. For large migration rates, however, we underestimate the true probability of rescue. This is because we ignore the temporal change of the fitness of rescue mutations. In particular, we underestimate the establishment probabilities of mutations that occur at the end of phase 1, just before the environment in deme 2 deteriorates. Our approximation ignores this change in environmental conditions in deme 2 and hence assumes that individuals carrying mutations that occurred during phase 1 will be counter-selected in deme 2, even during phase 2 when they are actually positively selected in that deme. This effect is negligible for small migration rates but can have considerable effect for large migration rates. Because our model underestimates the rescue chance for migration rates slightly larger than the swamping limit, this might also explain why we do not see an increase in the chance for evolutionary rescue for very large migration rates in simulations.

Importantly, the probability of survival for *m* → 0, as well as the optimal intermediate migration rate that maximizes the chance of rescue are correctly estimated by equation (2), at least for mutants with a large initial disadvantage *s* (Figs. S6A, S7A and S8). For small *s* and small *θ*, the temporal inhomogeneity in selection coefficients becomes more important, as mutations may take a long time to escape drift and eventually establish. This effect is weak for small migration rates, but with high migration rates, a relatively large number of mutants in deme 2 will be displaced to deme 1 where their establishment probability will increase (*e.g.* see fig. S4).

Another effect that we have ignored in our model is the increase in probability of rescue for high migration rates due to what Uecker et al. [2014] called “relaxed competition”. Density regulation in the non-deteriorated deme fills the habitat to carrying capacity at the end of each generation. For high migration rates, the non-deteriorated deme is strongly depleted and density regulation can increase the total number of mutants in a single generation (*e.g.* see figure S3 in the supplemental material to see the relaxed competition in a case without *de novo* mutations).

### When does intermediate migration favor rescue?

A key unresolved question for evolutionary rescue in structured populations is: when does gene flow facilitate evolutionary rescue as compared to two populations in isolation? Our model allows us to derive a condition for when intermediate migration helps chances of survival (as compared to no migration at all) by calculating when the derivative of 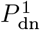 (that is, the probability of rescue due to *de novo* mutations during phase 1) with respect to *m* at *m* = 0 is positive. This is the case if (see supplemental material, Appendix B)

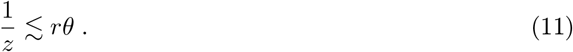

Thus, our model predicts that gene flow has a positive effect on evolutionary rescue if rescue mutations are strongly beneficial in the deteriorated environment (*z >* 0), respectively, if environmental change occurs slowly across demes (large *θ*), and/or if the new environment is very harsh (large *r*). The left hand side (11) simply quantifies the strength of positive selection. A larger selection coefficient of a rescue mutation increases the fitness gain of a mutant migrant that moves into the deteriorated deme. The right-hand side of condition (11) relates the strength of selection to the impact of demographic dynamics. Both *θ* and *r* influence the imbalance in population density between the two demes: the strength of stress, *r*, determines both the rapidity of decay of the population size in deme 1 as well as the equilibrium density of the population (see equation (9) and Fig. 1, as well as equation (S5) in Appendix A of the supplemental material). The length of an epoch *θ* determines the length of the period where deme 1 has a small population size relative to deme 2 such that gene flow is more likely to bring mutants into the deme where they are adapted to, rather than removing them from the deme where they can establish. Hence a long deterioration time or high stress extends the period where population size is low in deme 1 and large in deme 2, which is when gene flow has positive effects on rescue.

Figure 3 shows the comparison between analytical model and simulation for different combinations of parameters. In the first row 1*/z* ≥ *rθ*, and as predicted by theory we observe that simulations show a roughly constant probability of rescue over the range of the migration rate *m*. A small increase in the probability of rescue can be observed as *θ* increases (from left to right), in particular in the top-right plot (1*/z* = *rθ*). This increase is clearly observed in all subsequent rows (for higher *z*, top to bottom), confirming that condition (11) predicts when gene flow will facilitate evolutionary rescue.

**Figure 3:**
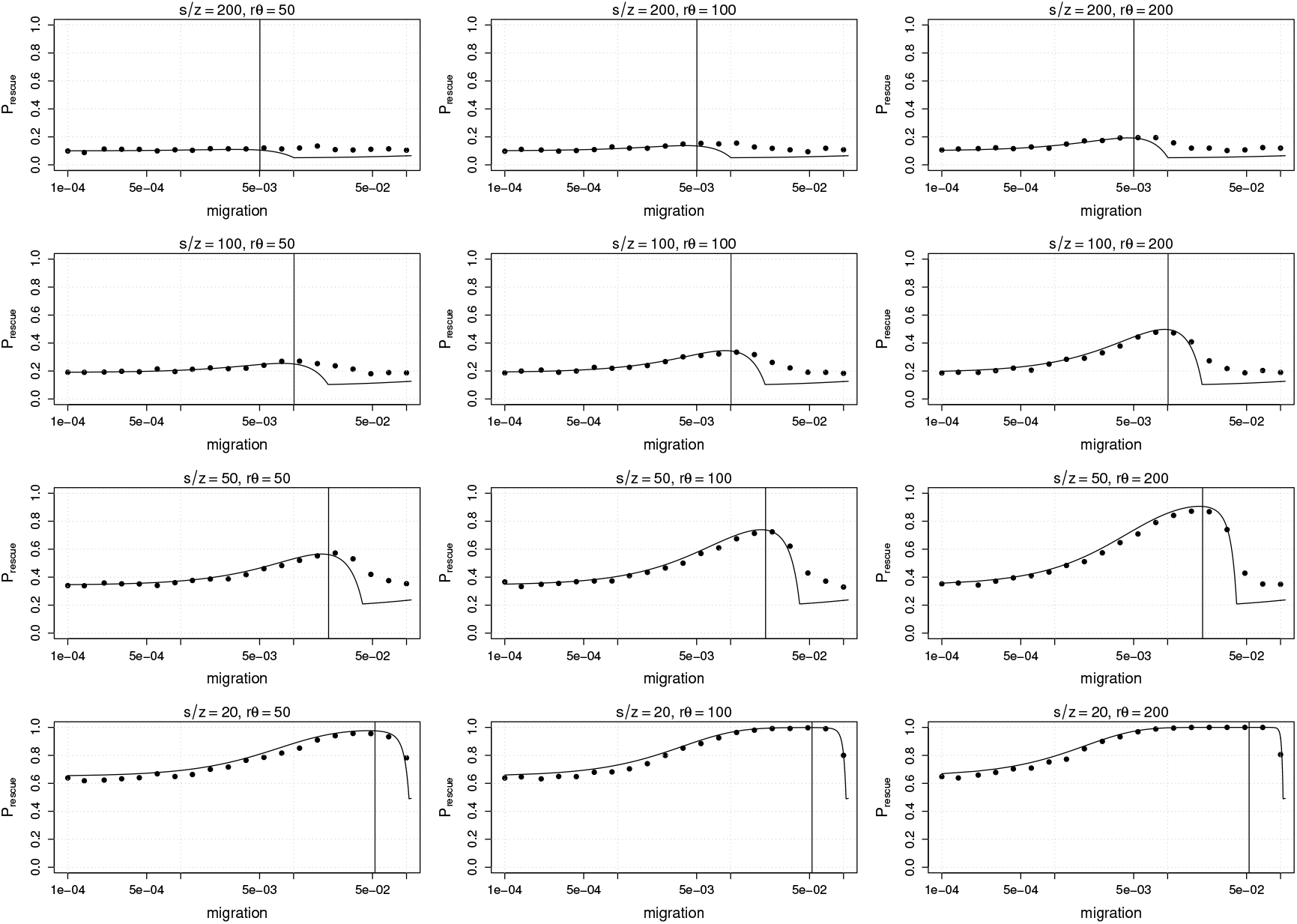
Evolutionary rescue for different combinations of parameters: first row *z* = 0.005, second row *z* = 0.01, third row *z* = 0.02, fourth row *z* = 0.05; left column *θ* = 500, center column *θ* = 1000, right column *θ* = 2000. In all figures, *r* = 0.1, *s* = 1.0. The vertical black line in each figure is the limit for swamping, *sz/*(*s z*). In the top two rows, we can see that passing from a situation where *s/z > rθ* to one where *s/z < rθ* makes the optimal migration rate more and more important. More extreme differences (*e.g.* third row, right column) yield a higher probability of evolutionary rescue at the optimal migration rate.

### Non-lethal rescue mutations

If we consider only *de novo* mutations, eq. (11) can be readily generalized to non-lethal mutations and becomes

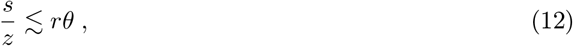

as is shown in the supplemental material (Appendix B). Note that this includes the condition (11) for lethal mutations as a special case if *s* = 1. If rescue mutations are sub-lethal or only slightly deleterious (*s <* 1), the range of parameters for which gene flow facilities evolutionary rescue increases. Migration is less detrimental because a mutant experiences a milder change in fitness when migrating from one deme to another. This is sensible as gene swamping is less likely if mutations are less deleterious in the environment to which they are not adapted [Bulmer, 1972, Lenormand, 2002, Tomasini and Peischl, 2018].

Unless the selective disadvantage *s* of rescue mutations is very large, rescue mutations will generally be present at low frequencies in the population before the deterioration of the environment. We thus need to account for the contribution of standing genetic variation to the probability of rescue (figure 4). We can see that the chances of survival from standing mutations are maximal in absence of migration (figure 4, also figure S3A). The reason is the following: a mutation in deme 1 at *t* = 0 will have higher chances of surviving compared to a mutation in deme 2, where it is counter-selected, that is, *p*^(1)^ *> p*^(2)^ for any combination of parameters. Further, because *p*^(1)^ is monotonically decreasing [Tomasini and Peischl, 2018], *P*_sgv_ tends to decrease with increasing migration rates (except if *s* is small and *m* is large, see Figure S3B). By adding the contribution of standing genetic variation (as calculated with (5)) the equivalent of condition (12) yields

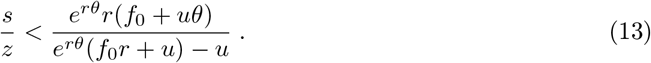

**Figure 4:**
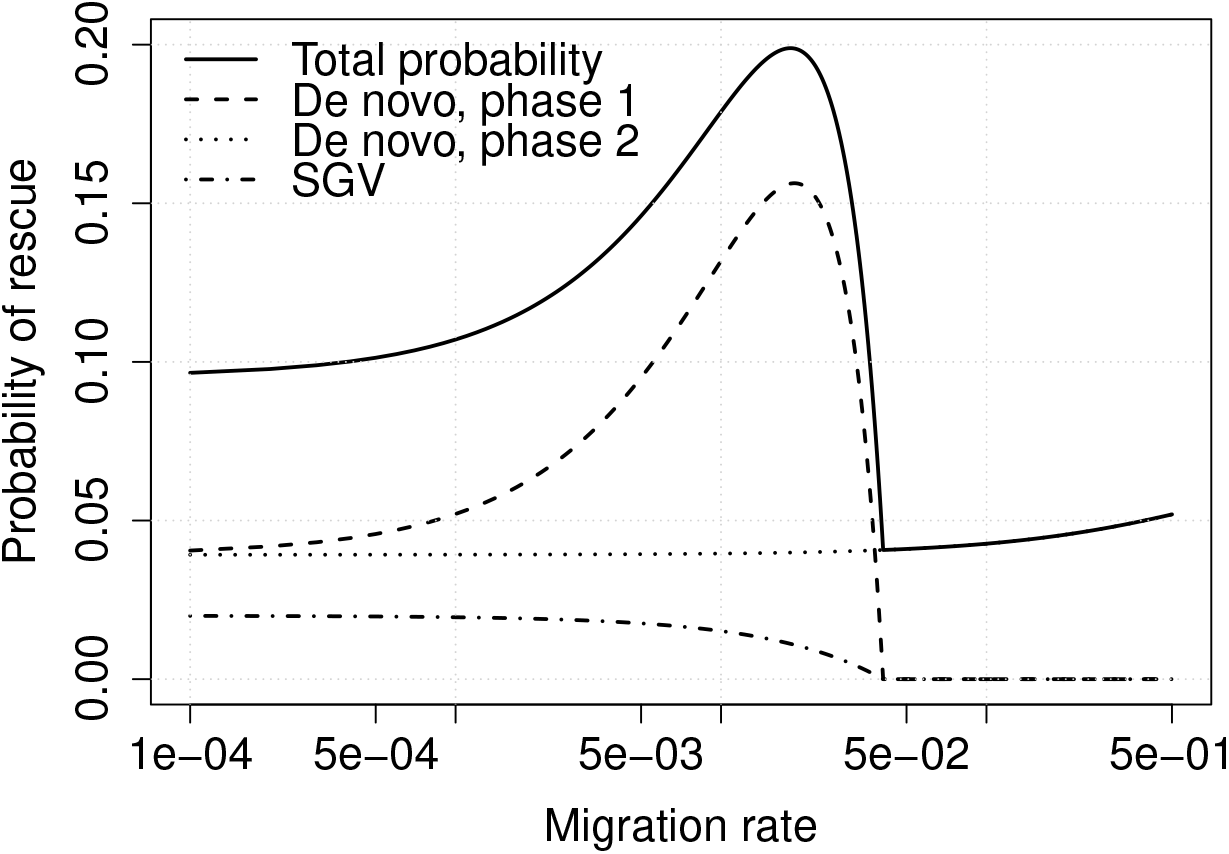
We show the total probability of rescue and its decomposition in terms of *de novo* mutations during phases 1 and 2, and standing genetic variation. Parameters are *z* = 0.02, *s* = 0.5, *r* = 0.5, *θ* = 500 and *f*_0_ = *u/s* (*i.e.* at mutation-selection equilibrium).

For *f*_0_ = 0, we recover equation (S11) in the supplemental material (Appendix B), which is in turn approximated to (12). When *f*_0_ increases, the right-hand part of (13) decreases, and gene flow loses importance. In fact, since *P*_sgv_ is monotonically decreasing with increasing migration rate *m*, standing genetic variation only matters for small to intermediate migration rates. Standing mutations will establish during phase 1 and are hence subject to gene swamping. Thus, if standing genetic variation is the predominant source of rescue mutations, gene flow is unlikely to have positive effects on rescue.

Figure S4 shows comparison between simulations and theoretical expectations for different values of *s* (with standing genetic variation). Our approximation is again very accurate for small value of *m*, whereas simulations and analytical approximations disagree for larger values of *m*. This disagreement is more pronounced for small values of *s*. This is due to new mutants that will spread so slowly that they will reach high frequencies only during phase 2, when both environments are deteriorated. The contribution of these mutants to the probability of rescue, however, is calculated through their probability of establishment in phase 1, which does not account for the temporal change in fitness of rescue mutations at time *θ*. The discontinuity between *p*^(*i*)^(*t < θ*) and *p*^(*i*)^(*t > θ*) causes our approximation to underestimate the probability of rescue, especially for large migration rates. Along these lines we also find that (13) is not accurate for small values of *s* (e.g., *s* = 0.1 in Figure S4). The analytical theory for standing genetic variation becomes accurate for sub-lethal mutations with a large selective disadvantage (e.g. Figure S8, *z* = 0.02, *s* = 0.5, *r* = 0.5, *θ* = 500, and *s/z* = 25 *<* 250 = *rθ*).

### Effects of the parameters of the model

Figure 5 illustrates the influence of various parameters on the probability of rescue. Increasing *z* has the main effect of increasing the probability of rescue, because a more beneficial mutation clearly has a larger chances of surviving (Figure 5A). At the same time, the optimal migration rate (when it exists) increases with increasing *z*. The reason is that the critical migration rate beyond which gene swamping occurs increases with increasing *z*: the condition for gene swamping is *m > sz/*(*s* − *z*) [Bulmer, 1972, Lenormand, 2002, Tomasini and Peischl, 2018]. For *z* ≪ 1, this reduces to *m* ≳ *z*, which thus allows establishment to occur for larger *m*. Decreasing the strength of environmental stress, *r*, leads to a higher overall probability of rescue because population sizes decline more slowly, leaving more time for rescue to occur (Figure 5B). The critical threshold at which swamping occurs remains unaffected, as it depends on the ratio between *z* and *m* only [Tomasini and Peischl, 2018]. Increasing *θ* extends the length of phase 1, which can increase the probability of rescue dramatically for intermediate migration rates but not for low or high migration rates (Figure 5C). For low migration rates, the length of phase 1 has very little impact since the two demes evolve almost independently. For strong migration, the length of phase 1 does not matter, because swamping prevents the establishment of rescue mutations during phase 1. Figure 5D shows that decreasing the deleterious effect of rescue mutations *s* has a similar effect on the probability of evolutionary rescue from *de novo* mutations as increasing *θ*. Decreasing *s* also affects the critical migration rate beyond which gene swamping occurs [Bulmer, 1972, Tomasini and Peischl, 2018], but this effect is rather weak. This can be seen if we rewrite the condition for gene swamping as *m > z/*(1 − *z/s*). In particular, if *z < s*, the effect of *s* becomes negligible.

**Figure 5:**
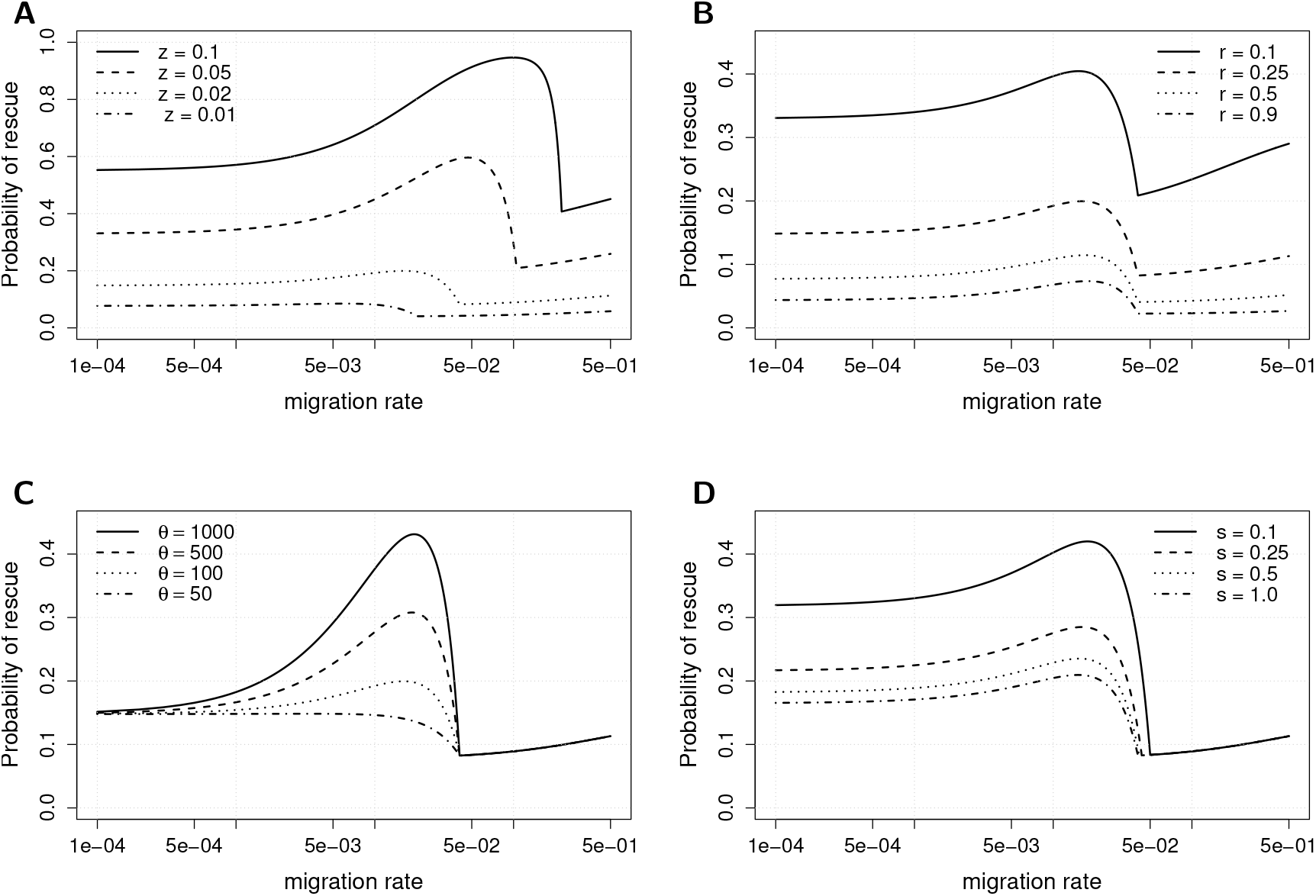
Total probability of rescue as a function of different parameters. When not otherwise stated in the legend, parameters are *z* = 0.02, *s* = 1.0, *r* = 0.25, *θ* = 200. (A) Variation with *r*, (B) variation with *θ*, (C) variation with *z*, (D) variation with *s* (and no standing genetic variation).

### Asymmetric carrying capacities and migration rates

We next consider the effect of asymmetric migration rates or asymmetric carrying capacities. For better comparison across models (see *e.g.* Barton et al. [2002]) and without loss of generality, we introduce two new parameters *ζ* and *β* that measure the degree of asymmetry:

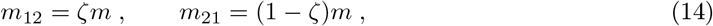

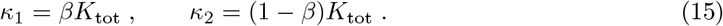

With these definitions, the model is symmetric with respect to migration rates if *ζ* = 0.5 and carrying capacities if *β* = 0.5. For *ζ <* 0.5, migration from deme 1 to deme 2 is smaller, while the opposite is true when *ζ >* 0.5. Figure 6A shows the probability of rescue as a function of *m* for different values of *ζ*. For *ζ* = 0.9, deme 2 receives many more migrants than it sends out, as compared to the symmetric model. The main effect of this asymmetry in migration is to decrease the total probability of rescue because rescue mutations are more likely to be removed from the deme to which they are adapted to as compared to the symmetric case. Further, gene swamping happens for lower values of *m* [Bulmer, 1972], thus reducing any beneficial effects of gene flow. The opposite is true for *ζ* = 0.1: wild-type individuals are removed at a smaller rate from the deme they are adapted to, which increases the chances of survival. At the same time, gene swamping occurs for larger values of *m* with respect to the symmetric case. The reduced effect of gene swamping with decreasing *ζ* also becomes apparent from the increase of the migration rate that maximizes the chance for evolutionary rescue. Figure S6A and S7A show comparison with simulations for *de novo* mutations and standing genetic variation with asymmetric migration rates.

**Figure 6:**
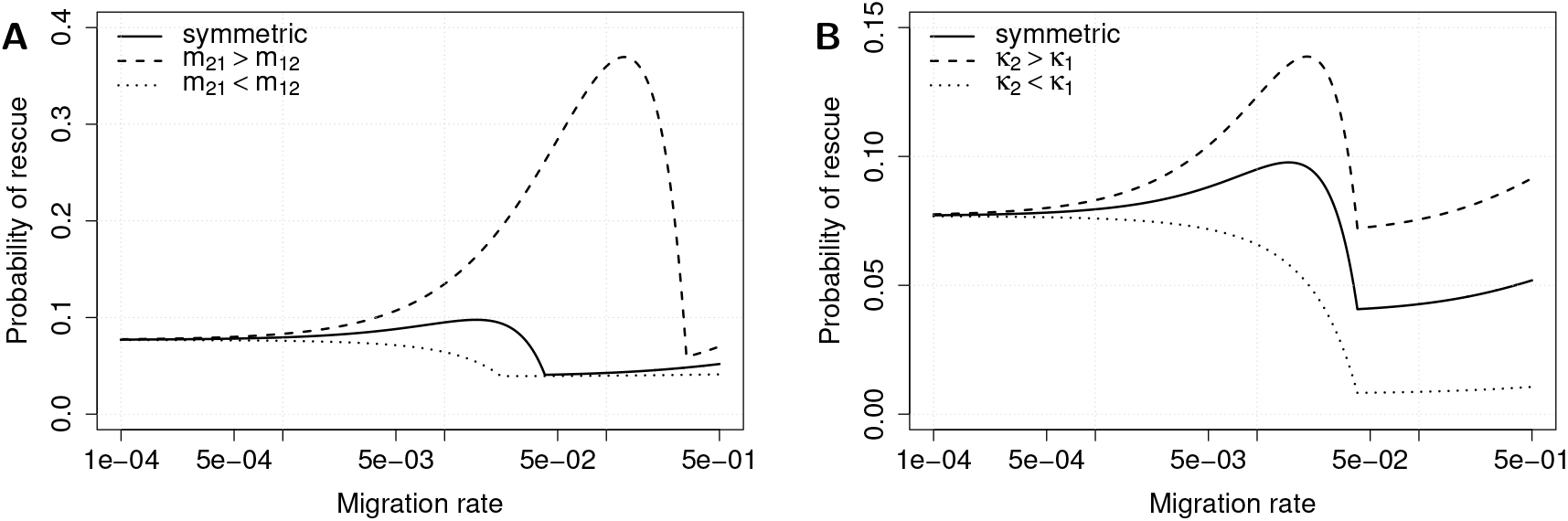
Probability of rescue as a function of migration for different sets of parameters and without standing genetic variation. *z* = 0.02, *s* = 0.5, *r* = 0.5, *θ* = 100, (A) *ζ* = 0.1, 0.5, 0.9, (B) *β* = 0.1, 0.5, 0.9.

We next keep migration rates symmetric, such that *m*_12_ = *m*_21_ = *m/*2, and investigate the effect of asymmetries in carrying capacities. Figure 6B shows the probability of rescue as a function of *m* for different *β*. We are going to call deme 2 “the reservoir”, as during phase 1 it is left untouched and it never goes extinct. We observe that a larger reservoir yields higher probability of rescue, and *vice versa*, when a reservoir is smaller the probability of rescue decreases. This is mainly due to *de novo* mutations during the second phase. Hence, chances of new mutants to establish increase because there are more wild-type individuals to start with at *t* = *θ*. When it exists, the optimal migration rate remains the same as in the symmetric model, even though it yields higher chances of survival for a larger reservoir. Figures S6B and S7B show comparison with simulations for *de novo* mutations and standing genetic variation with asymmetric carrying capacities. The condition for when gene flow facilitates evolutionary rescue from *de novo* mutations as compared to no migration becomes (see supplemental material, Appendix B)

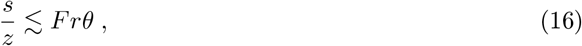

where

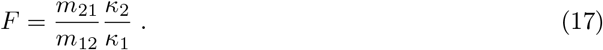

Condition (16) generalizes conditions (11) and (12) (it is also easy to generalize condition (13), as shown in the supplementary information, Appendix B, (S10)). This reflects the dynamics of a source-sink scenario. When deme 2 is large – the source is large – it sends many wild types to the sink, where new mutants could arise and prosper. The same happens if immigration in deme 1, *m*_21_, is large. In extreme cases, when *κ*_1_ *< m*_21_*κ*_2_, immigration in deme 1 causes overflow. This corresponds to a situation in which the population in a sink (in this case in deme 1) does not decline until the reservoir (deme 2) becomes deteriorated. On the other hand, since what matters most for ultimate rescue is the number of mutants, this high rate of migration also causes purifying selection in deme 1, not allowing any mutant to survive for long.

Figure S8 in the supplemental material (Appendix D) shows a comparison between theoretical expectations and simulations for asymmetric scenarios, revealing a good fit for small to intermediate migration rates.

### The role of density regulation

So far we have assumed that density regulation keeps the unperturbed deme at carrying capacity at all times. This requires sufficiently high local growth rates so that any reduction of the populations size due to emigration is immediately compensated by rapid growth within the unperturbed deme. This has the advantage that we do not need to model density regulation explicitly and is the same kind of density regulation as described in [Uecker et al., 2014]. We relax this assumption by assuming Beverton-Holt dynamics [Beverton and Holt, 1957] in the unperturbed deme: this means that the number of individuals *N_l_* of each type *l* (wild types or mutants, *l* ∈ {wt, m}) in the non-deteriorated deme in the next generation will follow

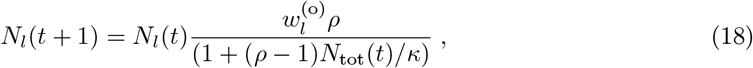

where *ρ* denotes the growth rate of the population, *N*_tot_(*t*) the total number of individuals in the deme, and 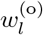 the fitness of individuals of type *l*. Differences between the two modes of density regulation are summarized in the supplemental material (Appendix C). We performed simulations of this model and compare the outcomes to the model with instantaneous growth (Figure 7). In all considered cases, the two modes of density regulation do not show any difference for low to intermediate migration rate. This is not surprising, as emigration affects the total number of individuals in the unperturbed deme only mildly, and even small values of *ρ* ensure that carrying capacity is maintained. For intermediate to large migration rates, however, the behavior can change dramatically (Figure 7). In particular, our simulations show that for large migration rates, the probability of rescue can be much lower if the growth rate *ρ* is small. To understand this behavior, let us first consider the case where population growth is instantaneous. The source population (unperturbed deme) is constantly losing individuals due to emigration into the sink population (perturbed deme). As a consequence, population growth will increase the absolute fitness of the remaining individuals in the source population [Tomasini and Peischl, 2018]. Thus selection in the unperturbed deme is less efficient as compared to the case without gene flow. The increase of the probability of rescue as *m* increases is due to relaxed competition and has been demonstrated formally in a two-deme model with source-sink dynamics [Tomasini and Peischl, 2018]. But if density regulation is logistic and growth rates are small, the advantage of relaxed competition disappears as emigration removes individuals more quickly than they can be reproduced. In this case we would expect that the probability of rescue starts to decline once the migration rate exceeds the critical value beyond which population growth can no longer maintain the population at carrying capacity. To calculate this critical migration rate, we approximate the net loss of individuals due to migration in deme 2 by solving

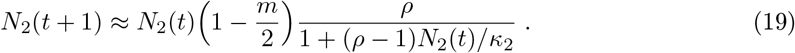

**Figure 7:**
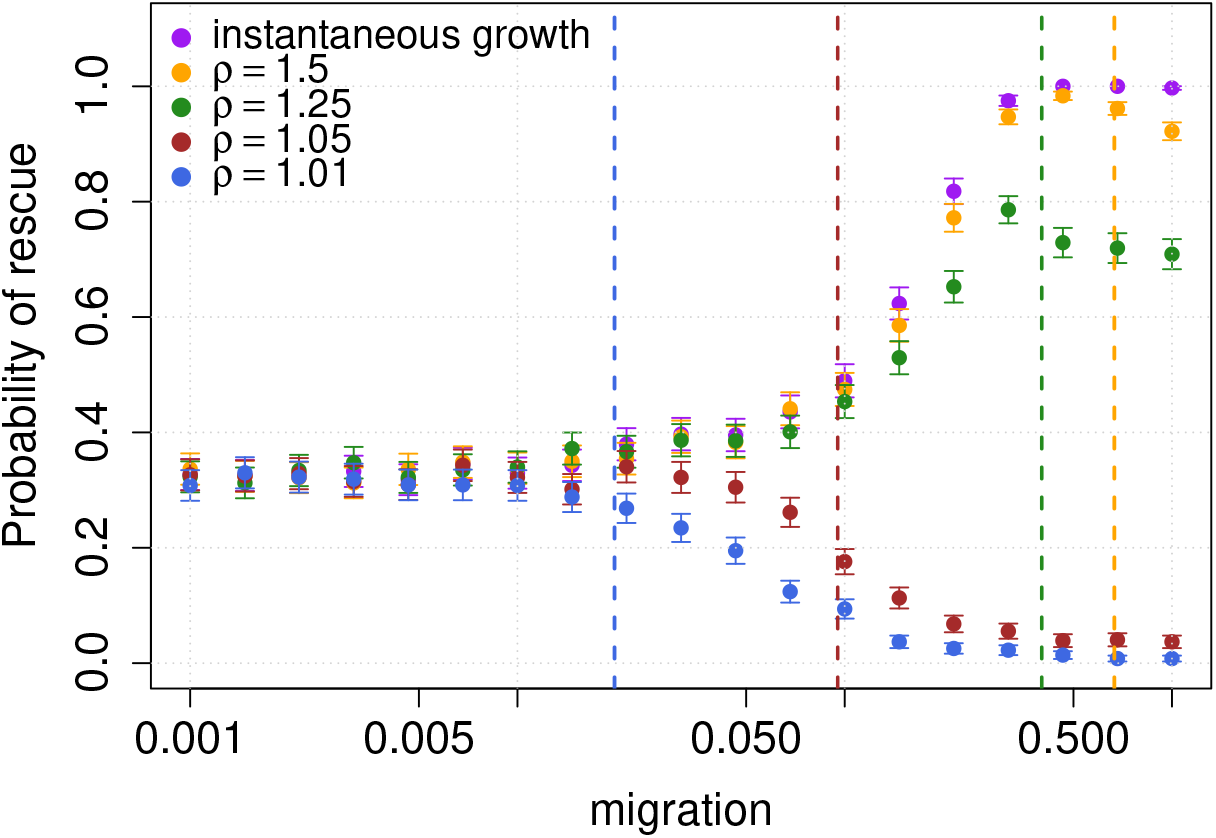
Comparison between different types of density selection for harsh changes over short periods. Here, *z* = 0.02, *s* = 0.1, *r* = 0.9 and *θ* = 100. The vertical lines show the critical migration rate for which equation (20) holds. Points and lines in blue refer to *ρ* = 1.01, in green *ρ* = 1.25, in orange to *ρ* = 1.5 and we show hard density regulation in purple.

Note that in this calculation we neglect the number of individuals coming from deme 1 and all the mutant individuals. The evolution of the individuals in deme 2 is calculated explicitly in the supplemental material (see Appendix C, equation (S14)). Now, extinction occurs when *N*_2_(*t*) = 0 for some *t >* 0. This happens when

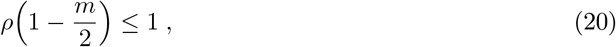

or when the product of the rate of growth and the rate of migration (loss) is smaller than 1. We should note that relation (20) is a conservative limit. As we do not take into account the presence of mutants, but only the net loss of wild-type individuals, this result does not account for the possibility of having a mutant establishing in the first generations after the deterioration event, as it is often the case [Peischl and Kirkpatrick, 2012]. The vertical lines in Figure 7 indicate this critical migration rates and confirm our intuitive explanation above.

Hence, density regulation can reduce the beneficial effects of gene flow if the growth rate *ρ* is not large enough such that the unperturbed deme does not remain at carrying capacity, and there is no relaxed competition. Even when there is the potential for relaxed competition in terms of *s*, *r* and *θ* (see [Uecker et al., 2014]), a slower growth rate lowers the chances of rescue for intermediate migration rates and higher (see figure 7). Ultimately, small growth rate *ρ* disrupts all effects due to migration and allows gene swamping to occur more readily. This is sensible, as low growth rate means that there will be fewer individuals in deme 2 and migration is mainly detrimental to the establishment of rescue mutations and also reduces the population size that can contribute to evolutionary rescue.

## Discussion

We studied a model for evolutionary rescue in a structured population using recent analytical results for establishment probabilities in structured populations [Tomasini and Peischl, 2018]. Our main result is an analytical prediction for the conditions under which gene flow facilitates evolutionary rescue in structured populations as compared to a population without gene flow. The potentially positive effect of gene flow on evolutionary rescue has been described previously both experimentally and theoretically; experimentally during adaptation to a gradient of salinity in a yeast meta-population [Gonzalez and Bell, 2013], mathematically in a model for evolutionary rescue in structured populations [Uecker et al., 2014], and via simulations of the evolution of treatment resistance in solid tumours [Waclaw et al., 2015]. These findings are in contrast to the fact that dispersal does generally not have a positive effect on (local) adaptation [Bulmer, 1972, Holt and Gomulkiewicz, 1997, Lenormand, 2002] in populations with more stable demographic scenarios, and the conditions for when gene flow facilitates survival in the face of drastic environmental change were previously not known. Our study fills this gap and provides surprisingly simple and intuitive conditions for when we expect positive effects of gene flow on survival via adaptation. Furthermore, our model allowed us to describe the interactions between density regulation, demographic dynamics and gene flow during adaptation to severe environmental stress.

We showed that the probability of evolutionary rescue from *de novo* mutations will be maximized for a migration rate *m >* 0 if *s/z < rθ*, where *r* describes the harshness of the new environment, *θ* the speed of environmental change, *s >* 0 is the cost of carrying a rescue mutation in the original environment (e.g., the cost of having a antibiotic mutation in the absence of antibiotics), and *z >* 0 is the selective advantage of a rescue mutation in harsh environments (e.g., the advantage of carrying an antibiotic resistance mutation in the presence of antibiotics). Thus, our model predicts that gene flow has a positive effect on evolutionary rescue if (i) rescue mutations are strongly beneficial/weakly deleterious in the deteriorated/original environment, respectively, if (ii) environmental change occurs slowly across demes (large *θ*), and/or if (iii) the new environment is very harsh (large *r*). We then extended this result to account for the effects of standing genetic variation, asymmetry in carrying capacities and the direction of gene flow between demes. Finally, we investigate the details of density regulation and find that they strongly affect whether gene flow will facilitate survival or not. In particular, if local growth rates in unperturbed demes are so low that carrying capacities cannot be maintained due to emigration of individuals, positive effects of gene flow diminish. The predictions that we derive from the model are corroborated by stochastic simulations.

Our results show that the main positive effect of gene flow is during phase 1, *i.e.* during the epoch in which only one deme is deteriorated. Gene flow from the unperturbed deme into the perturbed deme provides the raw material which can increase the chance of evolutionary rescue as compared to two populations without gene flow. This phenomenon has recently been formally studied in a two-deme model with divergent selection, where gene flow can be beneficial to the rate of establishment of locally adapted mutations [Tomasini and Peischl, 2018]. This is reflected in the equation *s/z < rθ*; the stronger the source-sink dynamics of the unperturbed and perturbed habitat (large *r*) and the longer these source-sink dynamics last (large *θ*), the more likely it is that gene flow is beneficial for evolutionary rescue. This effect is further amplified if carrying capacities or gene flow is asymmetric such that more individuals migrate from the unperturbed to the perturbed habitat (*F >* 1 in eq. (16)). Our model matches the results found by Uecker et al. [2014], in particular in the range where gene swamping does not occur (see Fig. S2 for a direct comparison).

We found that interactions between gene flow and density regulation play an important role. Ultimately, when the growth rate *ρ* of the wild type in deme 2 is large enough to compensate emigration to deme 1, the system remains in a source-sink scenario (see *e.g.* Gomulkiewicz et al. [1999]) and gene flow can be beneficial for evolutionary rescue. Furthermore, if the growth rate is very large, we observe relaxed competition (see also Uecker et al. [2014]) which can counter the negative effects of rescue mutations in the unperturbed habitat. If, however, gene flow depletes individuals too quickly in the unperturbed deme such that density regulation cannot replace these individuals, the positive effects of gene flow disappear (Figure 7).

It has been argued that standing genetic variation, along with initial population density, is the main factor determining the chances of evolutionary rescue [Gomulkiewicz and Holt, 1995, Barrett and Schluter, 2008, Agashe et al., 2011, Lachapelle and Bell, 2012, Ramsayer et al., 2013, Vander Wal et al., 2013]. While we find that this is the case in the absence of gene flow or if gene flow is very high, we also find that the contribution of de novo mutations can dwarf the contribution of standing variation for intermediate migration rates (see e.g., Figure 2). Also, we find that not only the initial size of the total population plays a major role, but also the variation in population densities across habitats (Figure 6).

The main short-coming of our approach is the inability to account correctly for the time-inhomogeneity of selective coefficients of wild-type and mutant individuals. This becomes critical for mutants arising just before the second deterioration event, as their probability of establishment will be closer to 2*z* than the approximation we used. This discrepancy increases with increasing migration rate (see eqs. (3) and (4)) and decreasing *s* (as slightly deleterious mutations are less likely to be purged before time *θ*). Hence, for slightly deleterious mutations our model underestimates the probability of rescue (see figure S4). It would be interesting to generalize our approach in such a way to account correctly for time-inhomogeneous selective coefficients, which could be achieved by fusing the approaches of Peischl and Kirkpatrick [2012] and Tomasini and Peischl [2018]. This is, however, a mathematically challenging endeavour and beyond the scope of this paper. Another interesting extension of our model would be to account for more than two demes. This would allow us to study different modes of dispersal, e.g., island models vs. stepping stone model, and could help to explain experimental findings that show that the mode of dispersal can strongly influence a population’s chance of survival [Bell and Gonzalez, 2011].

In our analysis, we assumed mutations that establish in isolation from other genetic events that may interfere with the process (*e.g.* clonal interference, [Gerrish and Lenski, 1998]). Therefore, we expect our results to hold in species reproducing sexually with strong recombination. In diploid individuals, the degree of dominance of rescue mutations may impact the evolutionary dynamics or rescue mutations. If mutations are co-dominant or partially recessive, our results can be carried over to diploid models by redefining our parameters *s* and *z* as the fitness effects of mutations in heterozygotes in the two environments. By excluding competition with concurrent mutations from our analysis, we expect this model to be less predictive for organisms reproducing with low recombination rates - or for mutations occurring in regions with low recombination rate. However, some of our results could still be valuable, as many of the effects that we described depend strongly on ecological aspects (such as carrying capacities, growth rate, migration rate) and evolutionary rescue focuses on relatively short periods such that co-segregation of multiple mutations seems unlikely.

Our approach could help improve understanding some of the results found in experimental setups (*e.g.* Bell and Gonzalez [2011]) and in theoretical investigations (*e.g.* Uecker et al. [2014]) about the effects of dispersal on the probability of evolutionary rescue. The simple and intuitive analytical predictions are imperative for our understanding of evolutionary rescue in structured populations and help us sharpen our intuition about the interactions of ecological and evolutionary process on short time-scales. A setup similar to the one proposed by Bell and Gonzalez [2011], with sub-populations of yeast exposed to a gradient of salt changing in time would be ideal to test our predictions.

## Supplemental material

Supplemental material for this paper can be found online: https://github.com/mtomasini/EvolutionaryRescue/blob/master/sm_biorxiv.pdf.

## Acknowledgements

We thank Mark Kirkpatrick, Sally Otto and Katie Peichel for stimulating discussions on this subject. We also thank Joachim Hermisson and Laurent Excoffier for helpful comments on the first manuscript. Version 5 of this preprint has been peer-reviewed and recommended by Peer Community In Evolutionary Biology (https://doi.org/10.24072/pci.evolbiol.100098). We gratefully acknowledge helpful comments from Claudia Bank, as well as three anonymous reviewers.

## Conflict of interest disclosure

The authors of this preprint declare that they have no financial conflict of interest with the content of this article. Stephan Peischl is one of the PCI Evol Biol recommenders.

## Notes

https://github.com/mtomasini/EvolutionaryRescue

### Competing Interest Statement

The authors have declared no competing interest.

### Summary of Updates

Minor layout tweaks.

## References

D. Agashe, J. J. Falk, and D. I. Bolnick. Effects of founding genetic variation on adaptation to a novel resource. Evolution: International Journal of Organic Evolution, 65(9):2481–2491, 2011.

M. V. Ashley, M. F. Willson, O. R. Pergams, D. J. O’Dowd, S. M. Gende, and J. S. Brown. Evolutionarily enlightened management. Biological Conservation, 111(2):115–123, 2003.

R. D. Barrett and D. Schluter. Adaptation from standing genetic variation. Trends in ecology & evolution, 23(1):38–44, 2008.

N. H. Barton, F. Depaulis, and A. M. Etheridge. Neutral evolution in spatially continuous populations. Theoretical population biology, 61(1):31–48, 2002.

G. Bell. Evolutionary rescue and the limits of adaptation. Phil. Trans. R. Soc. B, 368(1610): 20120080, 2013.

G. Bell. Evolutionary rescue. Annual Review of Ecology, Evolution, and Systematics, 48:605–627, 2017.

G. Bell and A. Gonzalez. Evolutionary rescue can prevent extinction following environmental change. Ecology letters, 12(9):942–948, 2009.

G. Bell and A. Gonzalez. Adaptation and evolutionary rescue in metapopulations experiencing environmental deterioration. Science, 332(6035):1327–1330, 2011.

R. J.H. Beverton and S. J. Holt. On the dynamics of exploited fish populations, volume 19 of 2. Ministry of Agriculture, Fisheries and Food, 1957.

L. M. Bono, C. L. Gensel, D. W. Pfennig, and C. L. Burch. Evolutionary rescue and the coexistence of generalist and specialist competitors: an experimental test. Proceedings of the Royal Society B: Biological Sciences, 282(1821):20151932, 2015.

E. C. Bourne, G. Bocedi, J. M. Travis, R. J. Pakeman, R. W. Brooker, and K. Schiffers. Between migration load and evolutionary rescue: dispersal, adaptation and the response of spatially structured populations to environmental change. Proceedings of the Royal Society B: Biological Sciences, 281(1778):20132795, 2014.

M. Bulmer. Multiple niche polymorphism. The American Naturalist, 106(948):254–257, 1972.

O. Carja and J. B. Plotkin. Evolutionary rescue through partly heritable phenotypic variability. Genetics, pages 977–988, 2019.

S. M. Carlson, C. J. Cunningham, and P. A. Westley. Evolutionary rescue in a changing world. Trends in Ecology & Evolution, 29(9):521–530, 2014.

C. Chevillon, M. Raymond, T. Guillemaud, T. Lenormand, and N. Pasteur. Population genetics of insecticide resistance in the mosquito culex pipiens. Biological Journal of the Linnean Society, 68(1-2):147–157, 1999.

L.-M. Chevin, R. Gallet, R. Gomulkiewicz, R. D. Holt, and S. Fellous. Phenotypic plasticity in evolutionary rescue experiments. Philosophical Transactions of the Royal Society B: Biological Sciences, 368(1610):20120089, 2013.

C. De Mazancourt, E. Johnson, and T. Barraclough. Biodiversity inhibits species’ evolutionary responses to changing environments. Ecology Letters, 11(4):380–388, 2008.

P. J. Gerrish and R. E. Lenski. The fate of competing beneficial mutations in an asexual population. Genetica, 102(0):127, Mar 1998. ISSN 1573-6857. doi: 10.1023/A:1017067816551. URL https://doi.org/10.1023/A:1017067816551.

J. H. Gillespie. Population genetics: a concise guide. The John Hopkins University Press, 2004.

R. Gomulkiewicz and R. D. Holt. When does evolution by natural selection prevent extinction? Evolution, 49(1):201–207, 1995.

R. Gomulkiewicz and R. G. Shaw. Evolutionary rescue beyond the models. Philosophical Transactions of the Royal Society B: Biological Sciences, 368(1610):20120093, 2013.

R. Gomulkiewicz, R. D. Holt, and M. Barfield. The effects of density dependence and immigration on local adaptation and niche evolution in a black-hole sink environment. Theoretical population biology, 55(3):283–296, 1999.

A. Gonzalez and G. Bell. Evolutionary rescue and adaptation to abrupt environmental change depends upon the history of stress. Philosophical Transactions of the Royal Society of London B: Biological Sciences, 368(1610), 2013. ISSN 0962-8436. doi: 10.1098/rstb.2012.0079. URL http://rstb.royalsocietypublishing.org/content/368/1610/20120079.

J. B. S. Haldane. A mathematical theory of natural and artificial selection, part v: Selection and mutation. Proc. Cambridge Phil. Soc., 23:838–844, 1927.

R. D. Holt. Population dynamics in two-patch environments: some anomalous consequences of an optimal habitat distribution. Theoretical population biology, 28(2):181–208, 1985.

R. D. Holt and R. Gomulkiewicz. How does immigration influence local adaptation? a reexamination of a familiar paradigm. The American Naturalist, 149(3):563–572, 1997.

D. Hughes and D. I. Andersson. Evolutionary trajectories to antibiotic resistance. Annual Review of Microbiology, 71:579–596, 2017.

M. Kirkpatrick and S. Peischl. Evolutionary rescue by beneficial mutations in environments that change in space and time. Philosophical Transactions of the Royal Society B: Biological Sciences, 368(1610):20120082, 2013.

J. Lachapelle and G. Bell. Evolutionary rescue of sexual and asexual populations in a deteriorating environment. Evolution: International Journal of Organic Evolution, 66(11):3508–3518, 2012.

T. Lenormand. Gene flow and the limits to natural selection. Trends in Ecology & Evolution, 17 (4):183–189, 2002.

S. Lion, V. A. Jansen, and T. Day. Evolution in structured populations: beyond the kin versus group debate. Trends in ecology & evolution, 26(4):193–201, 2011.

M. Lynch. Evolution and extinction in response to environ mental change. Biotic interactions and global change, pages 234–250, 1993.

B. H. Normark and S. Normark. Evolution and spread of antibiotic resistance. Journal of internal medicine, 252(2):91–106, 2002.

H. A. Orr and R. L. Unckless. The population genetics of evolutionary rescue. PLoS Genetics, 10 (8):e1004551, 2014.

M. M. Osmond and C. de Mazancourt. How competition affects evolutionary rescue. Philosophical Transactions of the Royal Society B: Biological Sciences, 368(1610):20120085, 2013.

S. Peischl and K. J. Gilbert. Evolution of dispersal can rescue populations from expansion load. bioRxiv, page 483883, 2018.

S. Peischl and M. Kirkpatrick. Establishment of new mutations in changing environments. Genetics, pages 895–906, 2012.

H. R. Pulliam. Sources, sinks, and population regulation. The American Naturalist, 132(5):652–661, 1988.

J. Ramsayer, O. Kaltz, and M. E. Hochberg. Evolutionary rescue in populations of pseudomonas fluorescens across an antibiotic gradient. Evolutionary applications, 6(4):608–616, 2013.

P. Samani and G. Bell. Adaptation of experimental yeast populations to stressful conditions in relation to population size. Journal of evolutionary biology, 23(4):791–796, 2010.

K. Schiffers, E. C. Bourne, S. Lavergne, W. Thuiller, and J. M. Travis. Limited evolutionary rescue of locally adapted populations facing climate change. Philosophical Transactions of the Royal Society B: Biological Sciences, 368(1610):20120083, 2013.

M. Tomasini and S. Peischl. Establishment of locally adapted mutations under divergent selection. Genetics, 209(3):885–895, 2018. doi: 10.1534/genetics.118.301104.

H. Uecker. Evolutionary rescue in randomly mating, selfing, and clonal populations. Evolution, 71 (4):845–858, 2017.

H. Uecker and J. Hermisson. The role of recombination in evolutionary rescue. Genetics, 202(2): 721–732, 2016.

H. Uecker, S. P. Otto, and J. Hermisson. Evolutionary rescue in structured populations. The American Naturalist, 183(1):E17–E35, 2014.

E. Vander Wal, D. Garant, M. Festa-Bianchet, and F. Pelletier. Evolutionary rescue in vertebrates: evidence, applications and uncertainty. Phil. Trans. R. Soc. B, 368(1610):20120090, 2013.

B. Waclaw, I. Bozic, M. E. Pittman, R. H. Hruban, B. Vogelstein, and M. A. Nowak. A spatial model predicts that dispersal and cell turnover limit intratumour heterogeneity. Nature, 525 (7568):261, 2015.

