## Supplemental material for "When does gene flow facilitate evolutionary rescue?"

### When does gene flow facilitate evolutionary rescue? – Supplemental Material

April 17, 2020

#### Appendix A: probability of rescue

We calculate the probability of rescue from beneficial mutations in a two-deme model in which habitats deteriorate over time. A mutation has selective coefficient $z > 0$ in a deteriorated region and coefficient $s > 0$ in a non-deteriorated region. We distinguish three different temporal phases: (phase 0) at $t < t_0$, both demes are not deteriorated; (phase 1) at $t_0$ we deteriorate deme 1; (phase 2) at time $t = t_0 + \theta$ we deteriorated deme 2.

To evaluate equations (5)–(8) (at end of phase 0 and during phase 1) we use the probabilities of establishment of mutations experiencing different selection pressure in each patch of a two-deme model [Tomasini and Peischl, 2018]. A mutation can arise in deme 1 or in deme 2, and it establishes with probabilities $p^{(1)}$ and $p^{(2)}$ respectively:

$$p^{(1)} = z(1 + \sigma - \Delta) - s\mu_{12} \; , \tag{S1}$$

$$p^{(2)} = z\mu_{21} - s(1 - \sigma + \Delta) \; , \tag{S2}$$

where

$$\sigma = \frac{z+s}{\lambda} \; , \qquad \mu_{ij} = \frac{2m_{ij}}{\lambda} \; , \qquad \Delta = \frac{\mu_{12} - \mu_{21}}{2} \; , \tag{S3}$$

and $\lambda = \sqrt{(m_{12}+m_{21})^2 + (z+s)^2 - 2(m_{12}-m_{21})(z+s)}$. The derivation is based on slightly super-critical branching processes and is valid for large populations with weak selection ($1/N < z \ll 1$). The method is valid for slightly super-critical branching processes (see Tomasini and Peischl [2018] for a full discussion of the validity of the model). Note that these equations do not account for the temporal in-homogeneity in selection coefficients at time $t = \theta$, and hence should be a good approximation if $\theta \gg 0$ and if $s \gg 0$.

### Probability of rescue for *de novo* mutations

In order to calculate formula (7) we need to solve equations (9) and (10). During phase 1 ($t < t_0+\theta$), $N_2(t) = \kappa$. Solving (9) yields

$$N_1(t) = e^{-(m_{12}+r)t}\left[\kappa_1 - \frac{m_{21}\kappa_2}{m_{12}+r}\right] + \frac{m_{21}\kappa_2}{m_{12}+r}\ . \tag{S4}$$

##### Equilibrium population in deme 1 increases with $m$

In the following we assume that $\theta$ is large. For large $t < \theta$, the exponential term of equation (S4) goes to zero. For the symmetric case ($m_{12} = m_{21} = m/2$) then, population in deme 1 reaches an equilibrium described by

$$N_1(t) \approx \frac{m\kappa_2}{m+2r}\ . \tag{S5}$$

Then, this is also the population of deme 1 at time $t = \theta$. This shows that the population in deme 1 increases when $m$ increases (see main text, discussion of figure 2). Note that this approximation is only valid for $\theta \gg 0$.

During phase 2, when both demes are deteriorated, $N_1(t)$ and $N_2(t)$ follow equation (10) with initial conditions $N_1(\theta)$ given by (S5) and $N_2(\theta) = \kappa_2$.

Equation (S5) does not only represent the population in deme 1 at time $t = \theta$, but also during most of phase 1, as the first term of the right-hand side of equation (S4) decreases exponentially. We can show that the same is true for the case with asymmetric migration or asymmetric carrying capacities. With asymmetric migration ($m_{12} = \zeta m$ and $m_{21} = (1-\zeta)m$, see main text, equations (14) and (15)), for large $t < \theta$, equation (S5) yields

$$N_1(t) \approx \frac{(1-\zeta)m\kappa_2}{\zeta m + r}\ . \tag{S6}$$

This increases with $\zeta$, and hence we deduce that the larger migration is from deme 2 to deme 1 ($\zeta$ increases) the larger $N_1(t)$ is over time, and the larger the probability of rescue.

For $m_{12} = m_{21} = m/2$ and asymmetric carrying capacities (from the main text, $\kappa_2 = (1-\beta)\kappa$), the equilibrium population in deme 1 is

$$N_1(t) \approx \frac{m(1-\beta)\kappa}{m+2r}\ , \tag{S7}$$

which increases when $\beta$ decreases (hence when $\kappa_1$ becomes smaller than $\kappa_2$).

The solutions $N_1(t)$ and $N_2(t)$ can be obtained straightforwardly for $t > t_0 + \theta$ but are very long and it does not bear any use to write them explicitly here. Plugging everything into (7) and (8), we obtain a straightforward analytical formula for the probability of rescue from *de novo* mutations during phase 1. All calculations can be easily carried out with software such as *Mathematica*.

#### Gene swamping does not allow for rescue in phase 1 for high $m$

In the main text we discuss how for high migration rates $m_{ij}$, establishment during phase 1 cannot occur because of a phenomenon called gene swamping. Gene swamping arises generally in two-deme models with divergent selection. Establishment probabilities during phase 1 (equations (S1)–(S3)) are 0 if $m > \frac{sz}{(s-z)}$ (in the case of symmetric migration, but the same can be calculated for asymmetric migration, see Tomasini and Peischl [2018]). This condition is equivalent to the gene swamping limit in deterministic models derived in Bulmer [1972] and Lenormand [2002]. Thus, rescue mutations cannot occur during phase 1 if the migration rate exceeds this limit.

### Appendix B: when does gene flow facilitate rescue?

We want to know for which set of parameters intermediate migration increases the chance for evolutionary rescue, as compared to no migration. This is equivalent to the set of parameters for which

$$\left.\frac{\partial P_{\text{res}}}{\partial m}\right|_{m=0} > 0\,. \tag{S9}$$

To do this, we use an approach similar to the one used in Tomasini and Peischl [2018] to find an approximated form of the probabilities of establishment (also see Appendix A). We first re-scale all parameters with respect to $z$ ($s = z\xi$, $m_{ij} = z\chi_{ij}$) and then linearize (2) with respect to $z$. Then, we take the derivative of the linearized form of $P_{\text{res}}$ and find its root. Switching back to the original variables, we find that condition (S9) is satisfied when

$$\frac{s}{z} < \frac{m_{21}}{m_{12}}\frac{\kappa_2}{\kappa_1}\cdot\frac{e^{r\theta}r(f_0+u\theta)}{e^{r\theta}(f_0 r+u)-u}\,. \tag{S10}$$

If we set $f_0 = 0$ (hence no standing genetic variation), we find that the condition reads

$$\frac{s}{z} < \frac{m_{21}}{m_{12}}\frac{\kappa_2}{\kappa_1}\cdot\frac{e^{r\theta}r\theta}{e^{r\theta}-1}\,. \tag{S11}$$

Because the function $xe^x/(e^x-1) \approx x$ if $x$ is large enough (approximately for $x \gtrsim 4$), we obtain equation (16). Note that conditions (11), (12) and (13) in the main text refer to the symmetric model ($m_{21}/m_{12} = \kappa_2/\kappa_1 = 1$, and with $s = 1$ in condition (11)), while here we derive the general result. In the main text we further define $F = m_{21}\kappa_2/m_{12}\kappa_1$.

#### Appendix C: the role of density regulation and local growth rates

Figure 7 shows examples for the probability of evolutionary rescue when the density of the non-deteriorated deme is regulated following a Beverton-Holt model of logistic growth [Beverton and Holt, 1957]:

$$N_2(t+1) = N_2(t)\frac{\rho}{(1+(\rho-1)N_{\text{tot}}(t)/\kappa)}\ . \tag{S12}$$

We find that the probability of rescue in this case can deviate strongly from the regime of instantaneous growth, where the population in the non-deteriorated deme always remains at carrying capacity. The latter should be a good approximation to the former if the growth rate $\rho$ is large enough relatively to the migration rate.

Here, we explore this intuition quantitatively. We calculate the loss of individuals from deme 2 during one generation, neglecting individuals coming in from deme 1. This works in particular for $t$ large, since deme 1 is almost depleted after a few generations and we can ignore the influx of immigrants from deme 1 into deme 2. Hence, we solve

$$N_2(t+1) = N_2(t)\left(1-\frac{m}{2}\right)\frac{\rho}{1+(\rho-1)N_2(t)/\kappa}\ , \tag{S13}$$

with initial condition $N_2(t=0)=\kappa$. We find

$$N_2(t) = \frac{\kappa(2-2\rho+m\rho)}{2-2\rho+2^t m\rho\left(\frac{1}{2\rho-m\rho}\right)^t}\ . \tag{S14}$$

Now, gene flow should be detrimental to evolutionary rescue if the interplay of $m$ (causing loss of individuals from deme 2) and $\rho$ (causing gain of individuals in deme 2) causes deme 2 to eventually go extinct. We find that $N_2(t)\to 0$, when $t\to\infty$, if

$$\rho\left(1-\frac{m}{2}\right)\leq 1\ . \tag{S15}$$

In particular, condition (S15) is very accurate for large $m$, as rescue for that range of migration is ensured exclusively by mutations arising during phase 2 (see figure 2). Figure 7 shows that this rule of thumb remains accurate over the whole range of $m$ when other kinds of density regulation are at play.

### Appendix D: figures

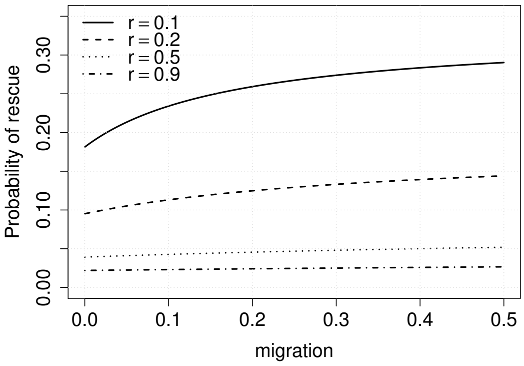


Figure S1: **Symmetric model:** contribution of mutations arising during phase 2 to evolutionary rescue for different $r$, $z = 0.02$, $s = 1$, $\theta = 200$. All curves increase with $m$.

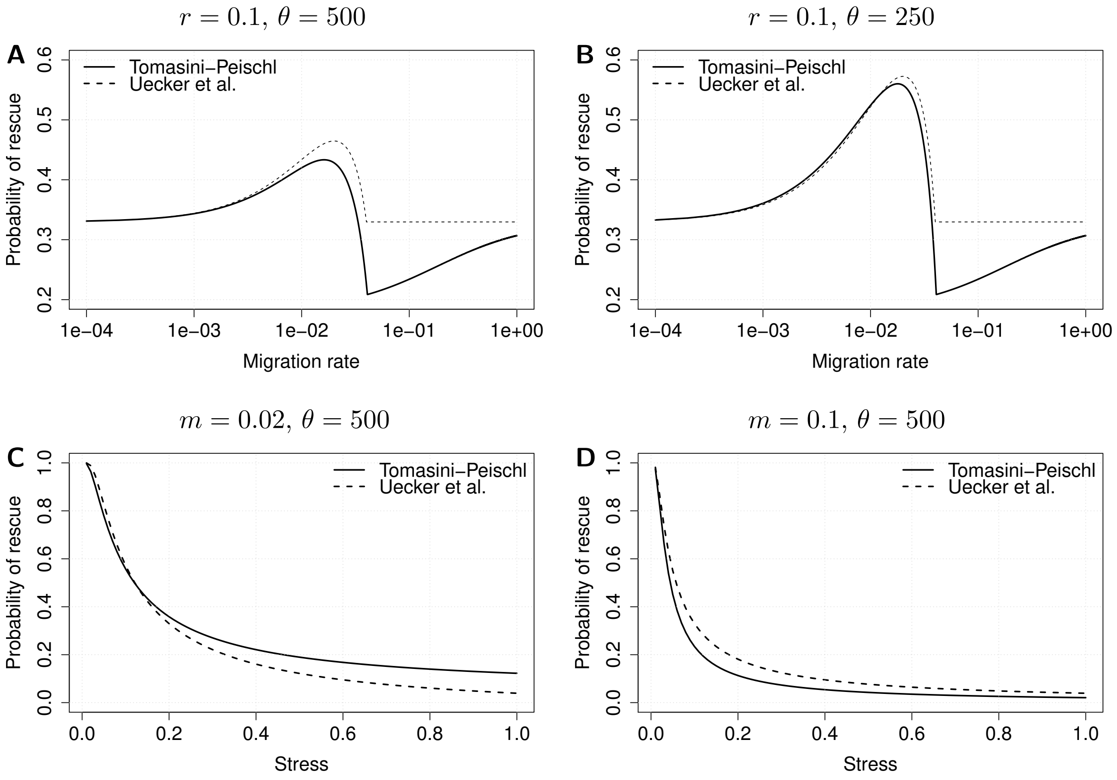


Figure S2: **Comparison:** (A and B) we plot approximation (2) in the main text VS. the approximation for two demes without standing genetic variation proposed by Uecker et al. [2014], with respect to the migration rate $m$, with $r = 0.1$ and (A) $\theta = 250$, (B) $\theta = 500$; (C and D) we plot the same comparison with respect to the stress $r$, with $\theta = 500$ and (C) $m = 0.02$ (corresponding to the ideal migration rate) and (D) $m = 0.1$. The two approximations are most similar for low $r$. Other parameters are $z = 0.02$, $s = 1.0$. Uecker et al. [2014] used a time-dependent process but did not model demes explicitly. Furthermore their solution is only for lethal mutations in the new environment ($s = 1$). We model both demes explicitly but do not take into account time-inhomogeneities, and our result is valid for $s < 1$. Our approach allows the derivation of general closed-form equations for the probability of rescue (while in Uecker et al. [2014] equations for every scenario need to be calculated separately). The two models yields very similar results for $m$ up to intermediate values.

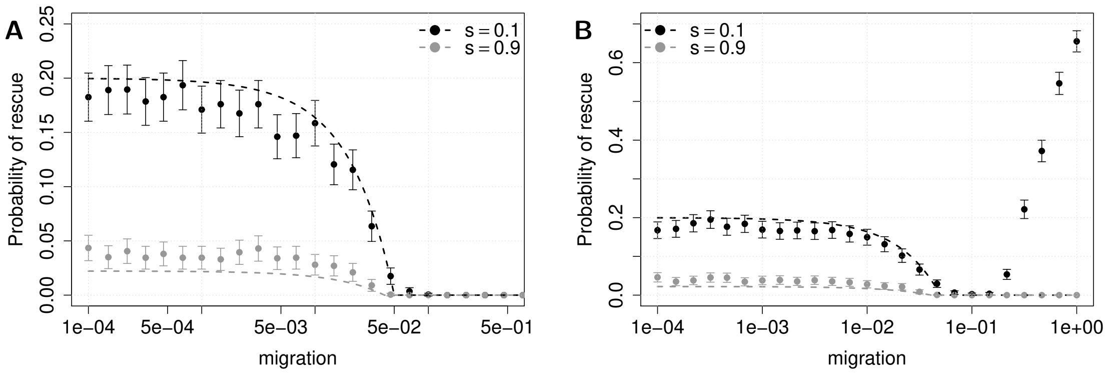


Figure S3: **Standing genetic variation:** contribution to evolutionary rescue by standing genetic variation in the symmetric model, simulations with analytical expectations (see equation (5). Parameters are $z = 0.02$, $\theta = 500$, $r = 0.3$. Black points show $s = 0.1$, gray points $s = 0.9$. (A) After density up-regulation, mutants are not replaced according to mutant frequencies preceding the regulation. (B) Mutants are replaced according to mutant frequencies. We can notice the effect of relaxed competition for mildly deleterious mutations (see main text).

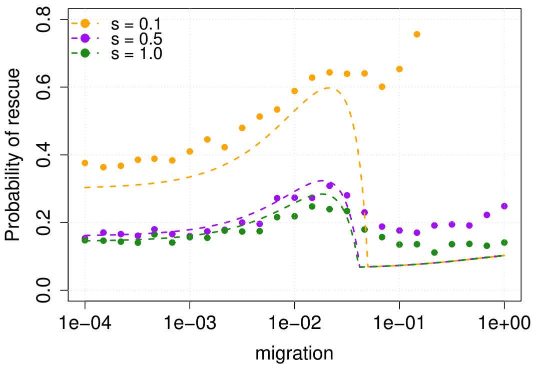


Figure S4: **Symmetric model:** evolutionary rescue as a function of $m$ for different selective coefficients $s$ (cost in the unperturbed deme). Comparison between theoretical calculations and simulations, for $z = 0.02$, $\theta = 500$ and $r = 0.3$, $s = 0.1$ (orange), $s = 0.5$ (purple), $s = 1.0$ (green). We observe that our model is unable to correctly account for mildly deleterious mutations (see orange line).

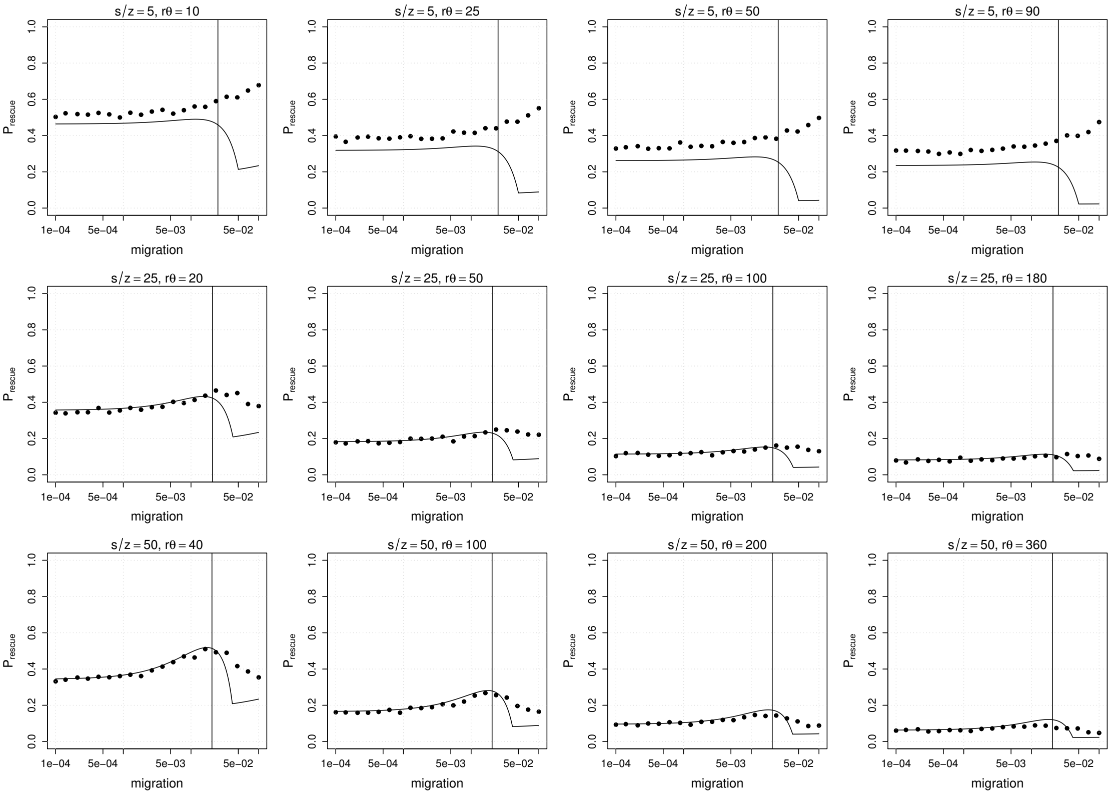


Figure S5: **Comparison between model and simulations for different combinations of parameters**: first row $s = 0.1$ and $\theta = 100$, second row $s = 0.5$ and $\theta = 200$, third row $s = 1.0$ and $\theta = 400$; left column $r = 0.1$, center left column $r = 0.25$, center right column $r = 0.5$, right column $r = 0.9$. In all figures, $z = 0.02$. The vertical black line in each figure is the limit for gene swamping, $sz/(s - z)$. In general, our approximation requires a very large $\theta$ to be precise. The values selected for $\theta$ in the first and second row are very low and we can observe that condition (12) in the main text is not clearly shown in simulations, in this case. Furthermore, the first row shows comparison between simulation and theory for very low $s$: we know that our model isn't able to correctly account for the time-inhomogeneity for such low values of $s$ (see also figure S4). Finally, choosing a very high value for $r$ (right column) yields very low probability of rescue, and simulations cannot clearly discern if gene flow facilitates rescue or not.

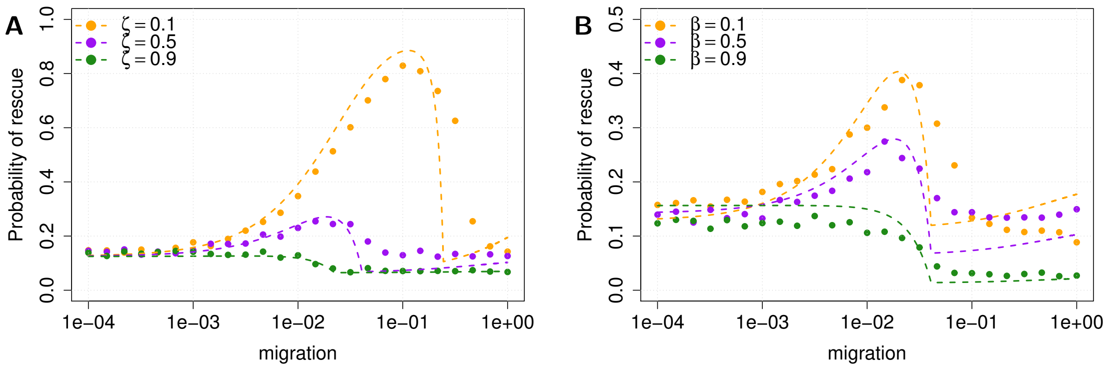


Figure S6: **Asymmetric models for lethal mutations:** comparison between theoretical calculations and simulations, for $z = 0.02$, $\theta = 500$, $r = 0.3$, $s = 1.0$. (A) Asymmetric migration rates. In orange, $\zeta = 0.1$, in purple $\zeta = 0.5$, in green $\zeta = 0.9$. (B) Asymmetric carrying capacities. In orange, $\beta = 0.1$, in purple $\beta = 0.5$, in green $\beta = 0.9$. We can see that at $m \to 0$ expectations for the model with asymmetric carrying capacities are not quite right, but still very close (see Appendix A).

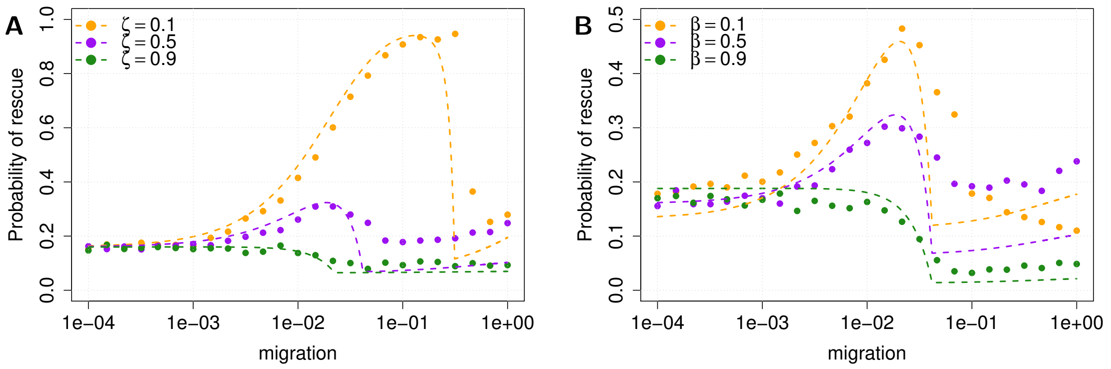


Figure S7: **Asymmetric models with standing genetic variation**: comparison between theoretical calculations and simulations, for $z = 0.02$, $\theta = 500$, $r = 0.3$, $s = 0.5$. (A) Asymmetric migration rates. In orange, $\zeta = 0.1$, in purple $\zeta = 0.5$, in green $\zeta = 0.9$. (B) Asymmetric carrying capacities. In orange, $\beta = 0.1$, in purple $\beta = 0.5$, in green $\beta = 0.9$. We can see that at $m \to 0$ expectations for the model with asymmetric carrying capacities are not quite right, but still very close (see Appendix A).

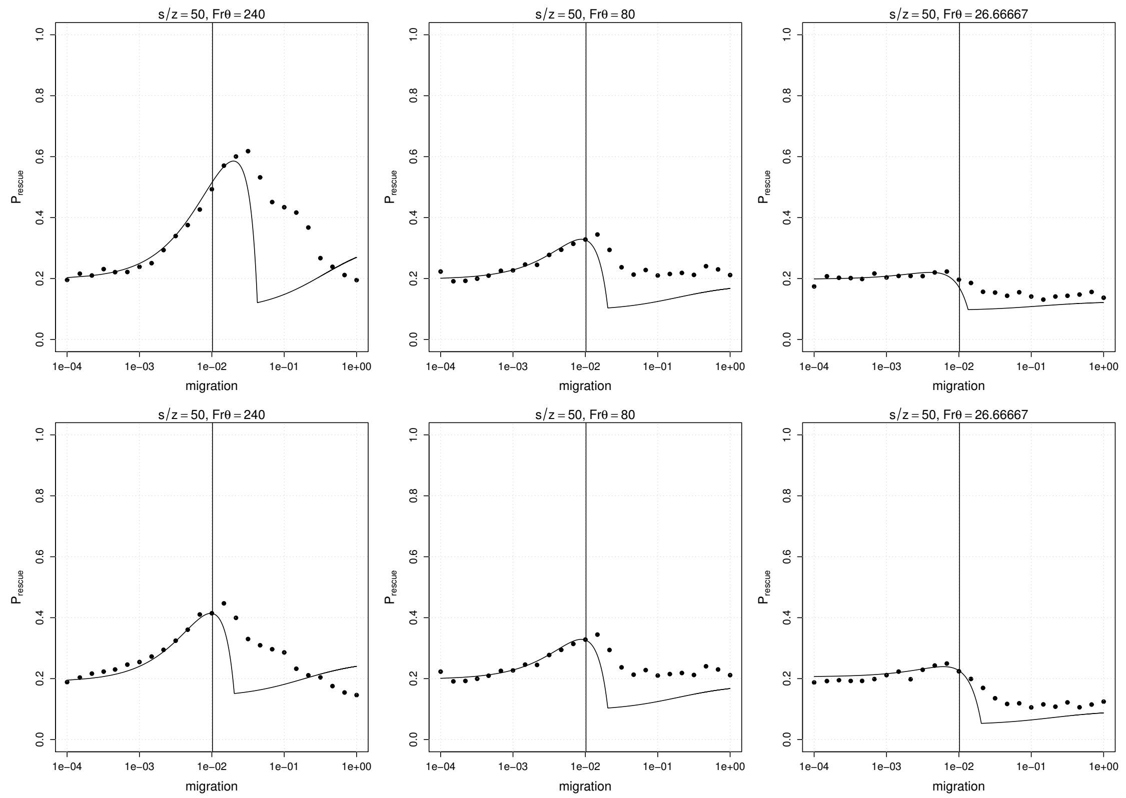


Figure S8: **Comparison between model and simulations for different combinations of parameters**: upper row is for asymmetric migration, lower row for asymmetric carrying capacities; left top panel has $\zeta = 0.25$, center top $\zeta = 0.5$ and right top $\zeta = 0.75$; left bottom has $\beta = 0.25$, center bottom $\beta = 0.5$ and right bottom $\beta = 0.75$. In all figures, $z = 0.01$, $s = 0.5$, $r = 0.1$, $\theta = 800$. The vertical black line in each figure is the limit for gene swamping, $sz/(s-z)$. This condition is calculated for a case with symmetric migration, thus it fails when $\zeta \neq 0.5$. Even in these scenarios, our model is able to predict reasonably well when migration facilitates evolutionary rescue. We also note that in simulations, while for the symmetric case (center top and bottom panels) $P_{\text{Rescue}}$ is the same for $m \to 0$ and $m \to 1$, this is not the case for both cases where there is an asymmetry in the system.

### References

R. J. H. Beverton and S. J. Holt. *On the dynamics of exploited fish populations*, volume 19 of *2*. Ministry of Agriculture, Fisheries and Food, 1957.

M. Bulmer. Multiple niche polymorphism. *The American Naturalist*, 106(948):254–257, 1972.

T. Lenormand. Gene flow and the limits to natural selection. *Trends in Ecology & Evolution*, 17 (4):183–189, 2002.

M. Tomasini and S. Peischl. Establishment of locally adapted mutations under divergent selection. *Genetics*, 209(3):885–895, 2018. doi: 10.1534/genetics.118.301104.

H. Uecker, S. P. Otto, and J. Hermisson. Evolutionary rescue in structured populations. *The American Naturalist*, 183(1):E17–E35, 2014.
